# Comparative single-cell transcriptomic analysis reveals putative differentiation drivers and potential origin of vertebrate retina

**DOI:** 10.1101/2023.12.03.569795

**Authors:** Xin Zeng, Fuki Gyoja, Yang Cui, Martin Loza, Takehiro G. Kusakabe, Kenta Nakai

## Abstract

Despite known single-cell expression profiles in vertebrate retinas, understanding of their developmental and evolutionary expression patterns among homologous cell classes remains limited. We examined and compared approximately 240, 000 retinal cells from four species and found significant similarities among homologous cell classes, indicating inherent regulatory patterns. To understand these shared patterns, we constructed gene regulatory networks for each developmental stage for three of these species. We identified 690 regulons governed by 530 regulators across three species, along with 10 common cell class-specific regulators and 16 preserved regulons. RNA velocity analysis pinpointed conserved putative driver genes and regulators to retinal cell differentiation in both mouse and zebrafish. Investigation of the origins of retinal cells by examining conserved expression patterns between vertebrate retinal cells and invertebrate *Ciona intestinalis* photoreceptor-related cells implied functional similarities in light transduction mechanisms. Our findings offer insights into the evolutionarily conserved regulatory frameworks and differentiation drivers of vertebrate retinal cells.

## INTRODUCTION

The retina acts as the primary sensory interface between the visual system and the external world and is crucial for vertebrate adaptation to varied ecological niches. Because of its structural complexity and accessibility, the retina is a pivotal model for studying the intricate functions of the central nervous system (CNS) (1, 2). For example, the presence of β-amyloid in both Alzheimer’s and age-related macular degeneration not only underscores the interconnected pathways of the retina and CNS, but also highlights the critical role of retina research in understanding and potentially treating CNS disorders (2). The vertebrate retina comprises one embryonic retinal progenitor cell (RPC) and seven mature cell classes that collaborate in visual information processing: rod and cone photoreceptors (PRs), bipolar cells (BCs), horizontal cell (HCs), amacrine cells (ACs), retinal ganglion cells (RGCs) and müller glia (MG) (3). Notably, certain developmental phases in the zebrafish retina feature ciliary marginal zone (CMZ) progenitors or retinal stem cells (RSCs) akin to RPCs (4). Unraveling the intricate regulatory mechanisms of the vertebrate retina has been challenging because of the high cellular heterogeneity and the various patterns of cell body distributions and connectivity (5).

The rapid advancement of single-cell RNA-seq (scRNA-seq) has enabled the detailed profiling of thousands of retinal cells, revealing molecular features of diverse cell classes in model organisms such as *Mus musculus* (mouse) (6), *Gallus gallus* (chicken) (7), and *Danio rerio* (zebrafish) (8). These studies have deepened our understanding of cellular identities within the vertebrate retina and laid the groundwork for comparative analyses (7, 9), emphasizing shared and unique transcriptomic landscapes across species. In addition, the field is moving beyond marker gene identification to exploring gene regulatory networks, offering deeper insights into regulatory mechanisms. Methods such as SCENIC (10), which predict gene regulatory networks based on co-expression and transcription factor binding information, have enabled a deeper insight into the regulatory mechanisms underlying cellular functions. While these approaches have been applied to the study of mouse retinal cells (11), a thorough understanding of the conserved regulatory networks in the vertebrate retinal cells is still an area awaiting further exploration.

Furthermore, identifying genes whose expression levels positively correlated with retinal cell differentiation, also known as putative driver genes, is crucial in understanding retinal development. Previous studies on retinal differentiation in mouse and zebrafish have primarily utilized methods based on transcriptional similarity (6, 8). However, these approaches often fall short in capturing the dynamic and complex cellular changes over time (12), thus limiting our ability to accurately identify cell class-specific putative driver genes. Recent advancements have introduced methods grounded in RNA velocity (13, 14), which model the direction and rate of gene expression by using the ratio of unspliced to spliced RNAs, thereby determining the direction and speed of individual cells. By combining RNA velocity with transcriptional similarity, these methods significantly enhance our ability to predict differentiation trajectories and identify putative driver genes crucial for development

(15). Identifying the conserved driver genes between mouse and zebrafish that underlie retinal differentiation is essential for advancing our understanding of the fundamental mechanisms governing retinal development across vertebrates.

In addition, considering the complexity of vertebrate retina, exploring corresponding cell types in simpler invertebrate organisms offers a feasible method to grasp the fundamental regulatory mechanisms of the vertebrate retina. *Ciona intestinalis* Type A (*Ciona*), one of the closest relatives of vertebrates, featuring a relatively simplistic nervous system and visual structures, is a fitting model to investigate the evolutionary origins of the vertebrate retina (16–18). Although a previous study suggested that dopamine-synthesizing cells that are located close to *Ciona* photoreceptors have some vertebrate AC characteristics (19), an in-depth exploration of shared expression patterns between vertebrate retina and *Ciona* PR-related cells is lacking.

To bridge these gaps, we systematically investigate the transcriptional profiles of vertebrate retinal cells from three representative species: mouse, chicken, and zebrafish (6–8), focusing on conserved gene regulatory networks (GRNs) and key driver genes for retinal cell differentiation. We also extend our analysis by comparing these profiles with human retinal organoid development (20) and the invertebrate *Ciona* (21) to trace the potential origins of retinal cells.

## MATERIAL AND METHODS

### Single-cell RNA-seq data of retinal cells from human organoid, mouse, chicken, and zebrafish

The FASTQ data from scRNA-seq retinal cells were retrieved from the GEO database for three species: mouse, chicken, and zebrafish (GSE118614, GSE159107, GSE122680). The FASTQ data of human retina organoid development data was collected from ArrayExpress under accession number E-MTAB-12714. Specifically, human retina organoid development data include sample from 13 hours (13h), 16h, 34h and 40h. The mouse dataset spans ten developmental stages: embryonic day 11 (E11), E12, E14, E16, E18, postnatal day 0 (P0), P2, P5, P8, and P14. Chicken data were collected on E12, E16, and E18. Zebrafish data were collected 24 hours post-fertilization (24hpf), 36hpf, 48hpf, 72hpf, and 14 days post-fertilization (14dpf).

CellRanger (Ver. 6.1.2) (22) was used to create original count matrices, using GRCh38, GRCm38, GRCg6a, and GRCz11 as the reference genomes. These raw count matrices were imported into R and analyzed using Seurat 4 package (23). Cells of low quality were identified and removed following specific criteria. For the data sets including cell type annotation (human, mouse and chicken cells at E16 and E18 stage), cells annotated as either non-retinal or low-quality were removed. We further confirmed the annotation by marker genes mentioned in the original papers (6, 7, 20). In the remaining cases (zebrafish and chicken cells at E12), cells expressing <200 genes and those with mitochondrial expression proportion >10% were excluded. For each stage without annotated information, filtered cells were normalized and scaled by *SCTransform* function, followed by principal component analysis (PCA) using top 3000 highly variable genes (HVGs). The top 20 principal components (PCs) were used for clustering based on a shared nearest-neighbor modularity optimization-based clustering algorithm with default parameters and range of resolutions from 0.05 to 0.8. Cells were annotated based on existing marker genes (Supplementary Table S1). To integrate samples from the same species, the filtered cells were integrated, normalized, and scaled using *SCTransform* function, followed by PCA using top 3000 HVGs. Next, Canek (24) was used to eliminate the batch effect by the top 20 PCs. Non-linear dimensionality reduction was conducted on the corrected data using UMAP.

### Transcriptional profile similarity analysis

Firstly, to reduce the noise from sequencing, we combined 10 cells within each cell class to construct pseudo-cells using all the retaining cells. Secondly, we collected 11, 000 one-to-one orthologs for the three species from biomaRt (25) and homologene (26) package using *getLDS* and *homologene* function, respectively. Then, we employed the orthologs to integrate the pseudo-cell count matrices, which was used as input to MetaNeighbor. Specifically, we utilized *run_MetaNeighbor_US* function to identify the similarity between cell-class pairs among retinal cell classes using about 350 HVGs generated by *get_variable_genes* function. In brief, MetaNeighbor uses HVGs to calculate Spearman correlations for gene expression across various cell classes in different species, establishing a measure of similarity. The strength of these correlations is then applied to compute the Area Under the Receiver Operating Characteristic, which determines how well these correlations can distinguish between similar and dissimilar cell types across species, thus quantifying the reliability of cell class conservation.

For unsupervised clustering, the integrated pseudo-cell count matrix was used as input to Seurat 4 package. The data were log normalized and scaled, followed by PCA using HVGs generated above. Next, Harmony (27) was used to eliminate the batch effect by the top 20 PCs. The corrected data was used for clustering based on a shared nearest-neighbor modularity optimization-based clustering algorithm with default parameters and a resolution of 0.01. Non-linear dimensionality reduction was conducted using UMAP.

### Ranking database development for chicken and zebrafish

We constructed the TF-motif ranking database for the two species according to the SCENIC tutorial instructions. First, TF lists for chicken and zebrafish were downloaded from the AnimalTFDB database (28). Then, the transcription factor binding motifs in both species were obtained from JASPAR (29), CISBP (30), DBcorrDB (31), and HOCOMOCO (32). Given that TF binding specificities are known to be conserved across species (33), the binding information from their human homologs was also used. Bind motif matrices for 80.6% (914 /1134) TFs in chicken and 44.3% (1128/2547) TFs in zebrafish were collected. The promoter sequences (upstream 5000bp and downstream 100bp from the transcription start site) used for ranking were downloaded from Ensembl (GRCg6a, GRCz11). Finally, promoter sequences were ranked.

### GRN reconstruction

We used pySCENIC (34) for the reconstruction of the gene regulatory network. In brief, we generated co-expression modules based on the scRNA-seq data using GRNBoost2. A ranking database was used to filter out indirect targets by comparing the preliminary gene set with the transcription factor binding predictions, retaining only those genes with strong evidence of direct regulation. The minimum number of genes in a module was set to 8, with all other settings at their default values. The ranking database for mouse was obtained from cisTarget database. Finally, the activity of each filtered module in each cell was estimated using AUCell (10). AUCell estimates the activity of specific gene sets in individual cells by calculating the area under the curve of the ranked gene expression distribution, assuming that if the genes in the set are highly active, they should appear earlier in the ranked list of gene expression. The inferred regulons were filtered by the regulon specificity scores and the expression specificity its regulator of a regulon. The regulon specificity scores are calculated by comparing the distribution of a regulon’s activity across all cells to its distribution within a specific cell class, converting the resulting divergence into a score that reflects how specific the regulon is to that cell class.

To visualize the interactions of cell-class specific GRNs at each stage from the three species at each stage, regulons were selected based on their scores and the expression specificity of their regulators. The filtered regulons and their regulators were then used to construct the GRNs using ggraph. Purple nodes represent regulators, while gray nodes indicate target genes. Interactions between regulators were represented by red edges, while other interactions were shown with gray edges. To highlight the number of genes regulated by each regulator, a centrality layout was used, with regulators controlling a larger number of genes tending to be positioned at the center of the network.

### Cross-species regulon comparison

We employed cell class-specific regulons for cross-species comparisons. First, gene names within these cell-specific regulons were transformed into their respective homologous gene names in other species, using the previously obtained set of 11, 000 one-to-one homologous genes. Subsequently, the AUCell package was used to assess the activity of each regulon within the cell datasets corresponding to each species. For evaluation, we incorporated data from mouse cells extracted at four developmental stages: E14, E18, P8, and P14. In the case of chicken, data was derived from the E16 and E18 stages, while for zebrafish, we utilized data across all five stages (24, 36, 48, 72 hpf, and 14 dpf). Finally, we computed the mean z-score of the cell-specific regulons within their homologous cell classes.

### RNA velocity analysis

First, to generate spliced/unspliced count matrices for samples from mouse and zebrafish, the BAM files generated by CellRanger (Ver. 6.1.2) were processed using velocyto (13) with the repeat mask annotation file downloaded from UCSC genome browser (https://genome.ucsc.edu/). Samples from the same species were then integrated into one loom file. For the zebrafish dataset, cells with unspliced read proportions lower than 7% were removed. As a result, unspliced reads accounted for 26% of the total in mouse and 14% in zebrafish. Within individual cell classes, these proportions ranged from 19% to 35% in mouse cells and from 9% to 17% in zebrafish cells (Supplementary Figures S1A and S1B).

The processed loom files were then used as input to Scanpy (35). Initially, genes with larger than 10 shared counts were retained. The top 2000 HVGs were then used for PCA, and the top 30 PCs were used for the calculation of first-/second-order moments for each cell, considering the 30 nearest neighbors. The moments were used to determine RNA velocities for differentiation using dynamics model. The transition matrix of cells was estimated by combining gene expression similarity and RNA velocities using CellRank (15). The weight for gene expression similarity was set to 0.2. Furthermore, the initial and terminal states were identified from the transition matrix. In addition, the lineage probabilities for each cell toward the terminal states of cell classes were computed. For each cell class, we identified the top 100 driver genes through their high correlation with lineage probabilities. For mouse, these genes had a median correlation of 0.14, p-value < 2.20 × 10^-12^ (Supplementary Table S2); for zebrafish, the median correlation was 0.27, p-value < 1.96 × 10^-112^ (Supplementary Table S3). To visualize the trends for driver genes during differentiation, we estimated the Diffusion (DPT) pseudotime for each cell (Supplementary Figures S1E and S1F). The estimated DPT pseudotime is consistent with stage information (Supplementary Figures S1C and S1D). Using this DPT pseudotime to order the cells, we then fit Generalized Additive Models to the gene expression data, weighting the contribution of each cell according to fate probabilities.

### GO analysis of top 100 cell class-specific driver genes

We employed the *compareCluster* function from the clusterProfiler (36) package for the analysis of driver genes identified in mouse and zebrafish. The analysis utilized the *enrichGO* function, targeting biological processes (BP) with org.Mm.eg.db and org.Dr.eg.db as the gene ontology database for mouse and zebrafish. We applied the Benjamini-Hochberg method for p-value adjustment, setting the q-value cutoff at 0.05 for significance. The top 5 significant GO terms for each cell class were plotted.

### Identification of DEGs in vertebrate retina and *Ciona* photoreceptors-related cells

We identified the vertebrate conserved differentially expressed genes (DEGs) for the vertebrate retina using the *FindAllMarkers* function by from Seurat in two datasets, applying criteria of an adjusted p-value <0.05 and setting ‘*only.positive = TRUE*’ to focus exclusively on positive markers:

1. In the integrated pseudo-cells dataset, generating pseudo-cell-DEGs.
2. In datasets from each species separately, we took intersections by the 11, 000 orthologs to generate ortholog-DEGs.

The final conserved DEGs were defined by retaining genes from pseudo-cell DEGs only if they were either in the ortholog-DEGs or HVGs from pseudo-cell-DEGs.

We collected scRNA-seq data of *Ciona* at late tailbud II and larval stage from the GEO database (GSE131155). The single-cell raw count matrices were generated using CellRanger (Ver. 6.1.2) with KH2013 (37) as the reference genome. Annotations for cellular identities were kindly provided by the original authors. We focused on cells related to PR, which led to the extraction of eight PR-related cell classes. The selected PR-related cells were normalized and scaled through *SCTransform* function. Subsequently, PCA was performed to identify the top 20 PCs, which were then used for batch correction by Canek. The corrected dimensions were further reduced in a non-linear fashion using UMAP. Marker genes were identified using the *FindAllMarkers* function, applying a threshold of an adjusted p-value < 0.01 and setting ‘*only.positive = TRUE*’ to focus exclusively on positive markers.

### Identification of Orthologous gene among mouse, chicken, zebrafish, and *Ciona*

Orthologous gene identification among the four species was conducted using OrthoFinder 2.5.4 (38). First, we downloaded the protein sequences of the mouse (GRCm38), chicken (GRCg6a), zebrafish (GRCz11), and *Ciona* (KH2013) from the Ensembl (39) and Ghost databases (37) (http://ghost.zool.kyoto-u.ac.jp/download_kh.html). To enhance accuracy and reduce computational complexity, only the longest transcript variant of a gene was used for multiple sequence alignments with MAFFT. Subsequently, orthogroups were estimated using the MCL graph clustering algorithm. 2, 724 single-copy orthogroups and 6, 374 orthogroups containing genes from mouse, chicken, zebrafish, and *Ciona* were identified. 5, 894 orthogroups that contained genes from all species with less than five copies in each species were retained for further analysis.

## RESULTS

### Overview of the study strategy

We collected and annotated scRNA-seq data from GEO database (Figure 1A). We initially focused on assessing the conservation of transcriptomes in vertebrate retinal cells across different species (Figure 1B). Subsequently, we applied SCENIC (10) to discern cell-class-specific regulatory networks within each species (Figure 1C). We thus identified conserved regulators and regulons across vertebrates, elucidating evolutionarily preserved regulatory frameworks. In addition, we utilized RNA velocity analysis to identify conserved driver genes that propel retinal cell differentiation in both mouse and zebrafish (Figure 1D) and extracted the regulators that drive differentiation by intersecting regulatory networks (Figure 1E). Finally, we collected scRNA-seq data from *Ciona* (Figure 1F), identified shared DEGs between vertebrate retinal cells and *Ciona* photoreceptor-related cells (Figure 1G), and inferred the potential origin of vertebrate retinal cells.

**Figure 1.**
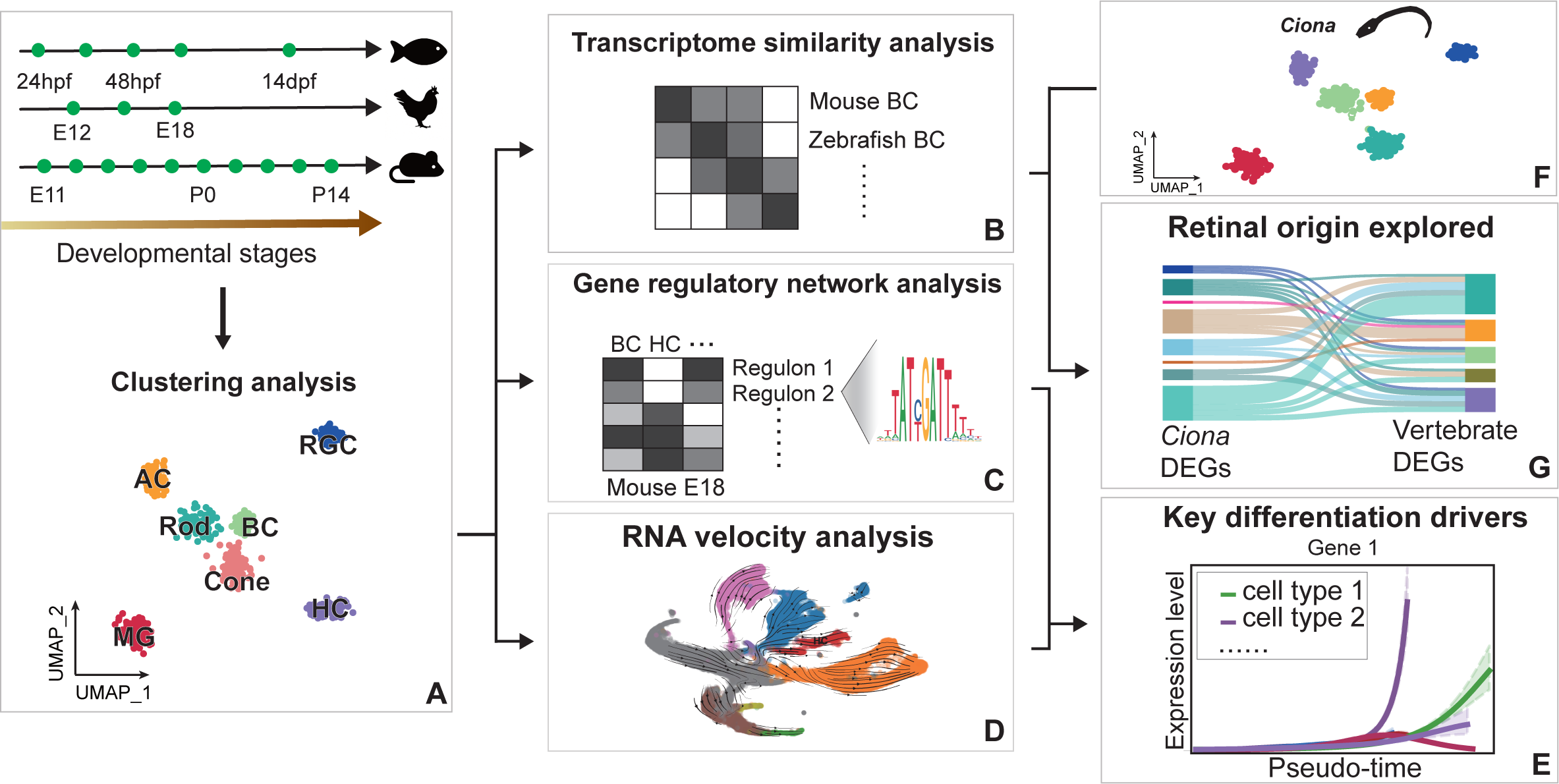
Workflow illustrating the transcriptional analysis of vertebrate retinal cells and their comparison with the invertebrate species *Ciona*. **A** scRNA-seq data from various stages of three representative species are collected and annotated. **B** Transcriptome similarity analysis showing transcriptome conservation among homologous retinal cells in vertebrates. **C** GRN analysis indicating cell class-specific regulatory networks and evolutionarily preserved regulatory frameworks within each vertebrate species. **D** RNA velocity-based analysis revealing conserved driver genes responsible for retinal cell differentiation in mouse and zebrafish. **E** Extraction of key differentiation-driving regulators through intersecting regulatory networks and RNA velocities. **F** scRNA-seq data from *Ciona* are collected. **G** Identification of genes with shared expression between vertebrate retinal cells and PR-related cells in *Ciona*, implying a potential ancestral link in their origin.

### Conserved expression patterns in vertebrate retinal cell landscapes

We downloaded scRNA-seq data of retinal cells for the three species from the GEO database (see Methods). Across the four species, we collected data obtained at 18 different time points. In total, about 240, 000 cells were retained for subsequent analysis, including 9704 human organoid cells (Supplementary Figures S2A), 100, 506 mouse cells (Figure 2A), 86, 056 chicken cells (Figure 2B), and 49, 311 zebrafish cells (Figure 2C). The data from all three species comprised seven mature cell classes distributed over stages (Supplementary Figures S2B-2E).

**Figure 2.**
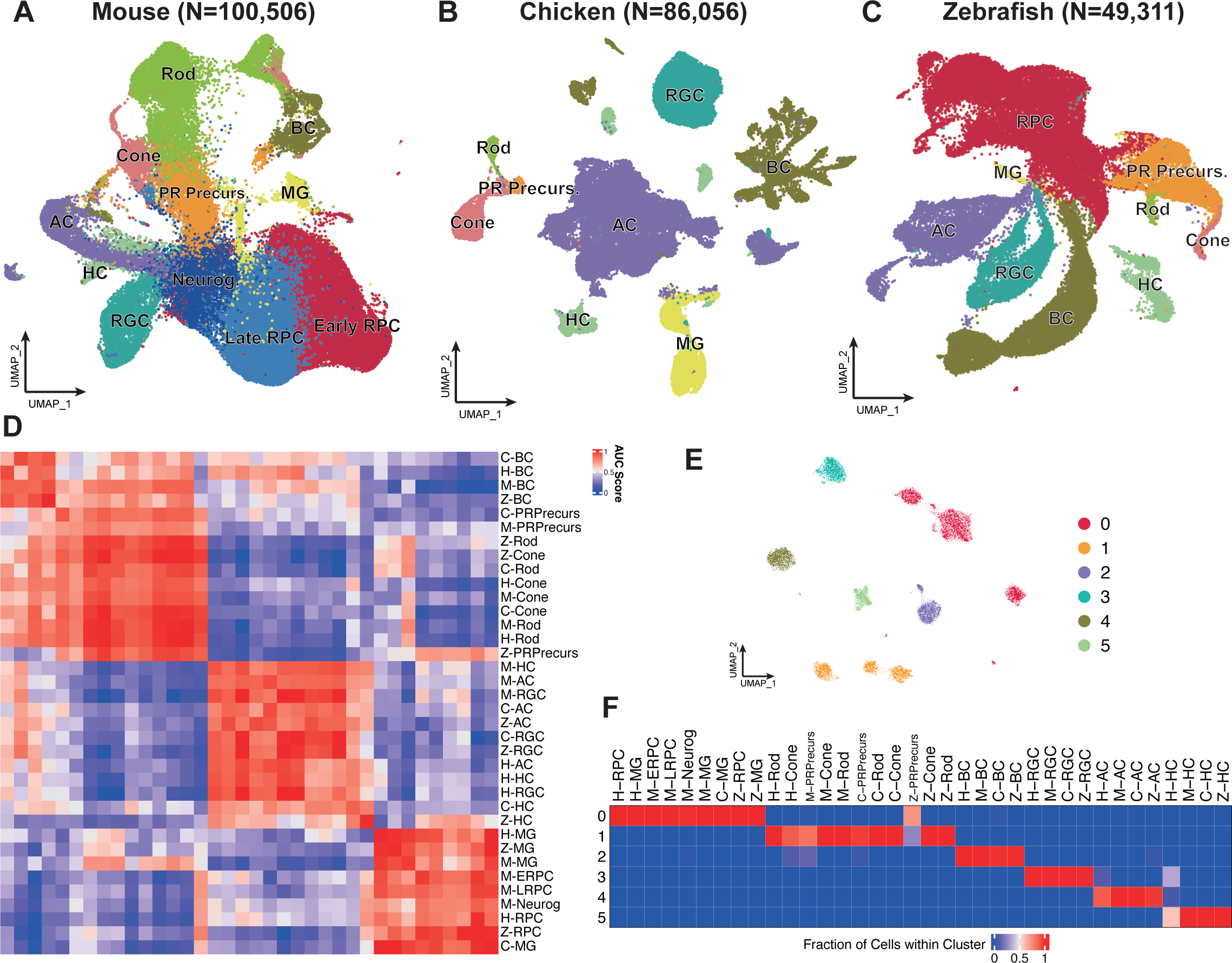
Comparative single-cell transcriptomic landscapes of retinal cells in mouse, chicken, and zebrafish. **A-C** UMAP plots of retinal scRNA-seq data from mouse (**A**, N = 100, 506), chicken (**B**, N = 86, 056), and zebrafish (**C**, N = 49, 311). **D** Heatmap of AUROC scores computed using MetaNeighbor, indicating similarity among cell classes based on orthologous gene expression from mouse, chicken, and zebrafish. In the heatmap, cell classes prefixes denote their origin: ‘H-’ for human, ‘M-’ for mouse, ‘C-’ for chicken, and ‘Z-’ for zebrafish. Red indicates high similarity, blue indicates low similarity. **E** Unsupervised clustering of pseudo-cells into six clusters displayed in UMAP. **F** Heatmap displaying the proportion of each cell class within the 6 clusters identified in **E**. Red indicates a high proportion; blue indicates a low proportion.

To evaluate transcriptome conservation across species, we compared cell similarities among the three species included in this study by first constructing pseudo-cells. This involved combining 10 cells within each cell class to reduce the noise from sequencing, using all the retained cells (see Methods). The heatmap (Figure 2D) revealed that the overall gene expression patterns in vertebrate retinal homologous cell classes were conserved, indicating that individual cell class displayed less variation in their transcriptome profiles during evolution than different cell classes within the same species. All MG shared similar expression patterns with RPCs, indicating their ability to differentiate into other retinal cell classes (40). Cones and rods shared more similar transcriptional profiles, which is consistent with their common role in converting light signals to neural signals (41). Additionally, ACs, HCs, and RGCs demonstrated closer relationships than other cell classes, indicating they share similar expression patterns. Unsupervised clustering of integrated pseudo-cell data revealed six major clusters (Figure 2E). Each cell class was confined to a specific cluster, and almost all the homologous cell classes were grouped into one cluster, indicating the similarity of expression profiles among homologous cell classes (Figure 2F).

### Identification and reconstruction of gene regulatory networks (GRNs) in vertebrate retina cells across species

To construct the GRNs for cells from the three species, we first built the TF-motif ranking database for chicken and zebrafish. Then, we estimated the GRNs for cell classes from each developmental stage from each species using the pySCENIC pipeline (34) (see Methods). Approximately 2700 regulons were inferred from the 18 datasets. An active regulon is defined as a collection of genes regulated by the same transcription factor (regulator) that are highly expressed early in the ranked list of gene expression in individual cells. The regulator of an active regulon was termed an ‘active regulator’. We calculated the regulon specificity scores for each regulon and extracted the top 20 cell class-specific regulons. We then removed low-quality regulons whose regulators were not specifically expressed in one cell class. Ultimately, 690 cell class-specific regulons were retained for further analysis (Figure 3A). We identified numerous regulators in various cell classes across different stages (Supplementary Figures S3A–S3G, S4A, S5A–S5D). Importantly, the regulators identified in mouse align with those discovered in a previous study (11), despite the study utilized a different method to infer gene regulatory networks. We identified known and novel retinal cell regulators across stages in the three species (Supplementary Tables S4-S6). Specifically, in E18 stage of mouse (Figure 3B), we identified well-established cell class-specific regulator such as Sox9 in late RPCs (42) and Gbx2 in a subclass of GABAergic ACs (43). Additionally, we identified regulators that have not previously been reported, such as Pitx1 and Pou3f1 in glycine ACs, and Zfp444 and Lcorl in PRs. Similarly, at the E18 stage in chicken (Figure 3C), we identified known regulators such as PAX6 in ACs (44) and THRB in cones (45), as well as previously unreported regulators such as LHX3 in BCs and HMX3 in HCs. At the 72 hpf stage of zebrafish (Figure 3D), we observed validated regulators such as sox9b in RPCs (46) and lhx1a in HCs (47). Furthermore, we identified previously unreported regulators such as hmx3a in HCs.

**Figure 3.**
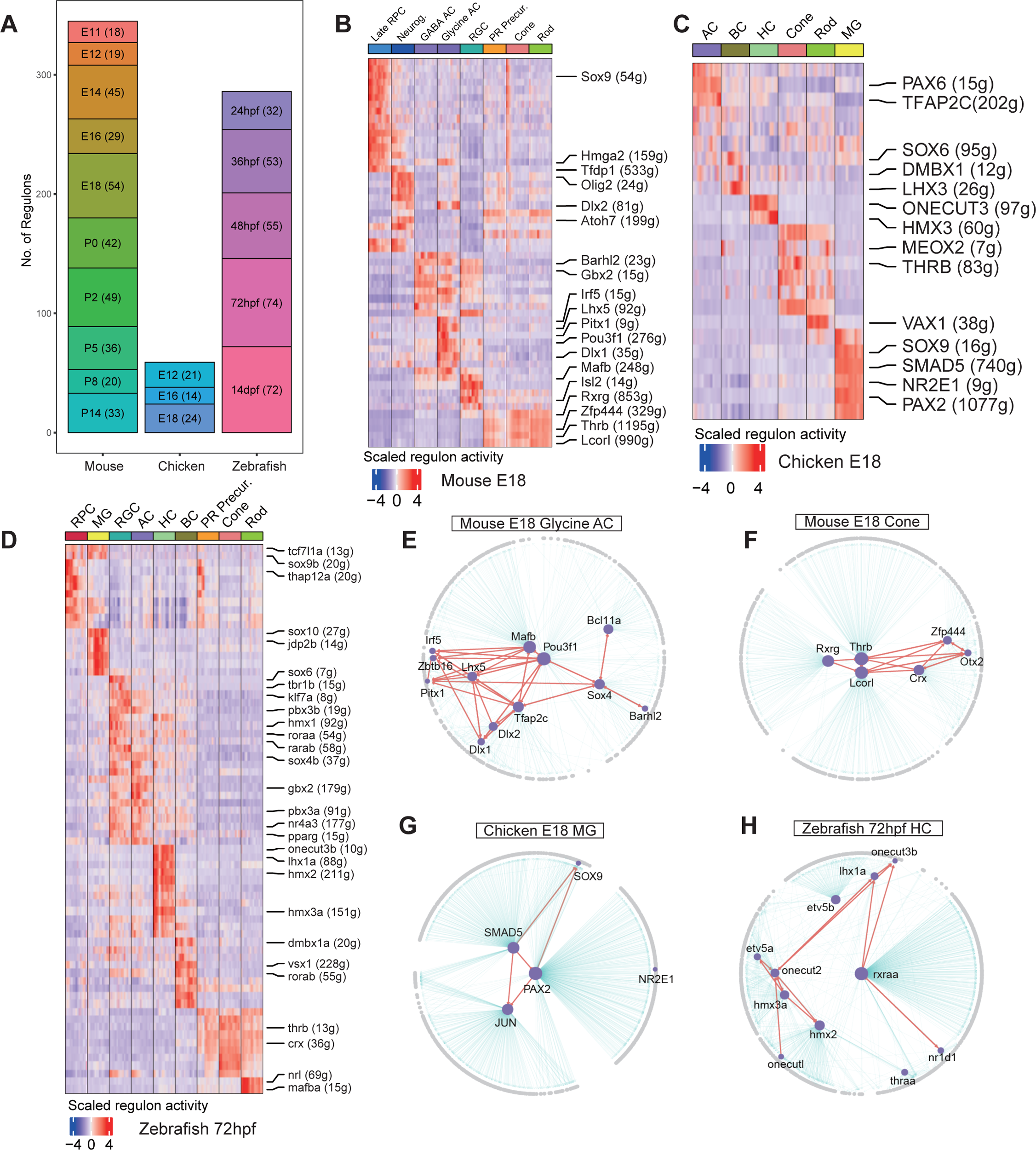
Gene regulatory network analysis in vertebrate retinal cells. **A** Number of identified and filtered regulons from scRNA-seq data across different stages in mouse, chicken, and zebrafish. Bar represents the number of regulons that have been filtered and identified from scRNA-seq data at distinct stages in the respective species. **B–D** Heatmaps of scaled regulon activity across different species and stages, including (**B**) mouse at E18, (**C**) chicken at E18, and (**D**) zebrafish at 72hpf. Each row represents a distinct regulon and each column denotes a specific cell class. Representative regulons for each cell class are highlighted in the right panel. Each regulon is named following the format “TF (N)”, where “TF” represents the regulator and “N” denotes the number of target genes in that regulon. Red indicates high activity, blue indicates low activity. **E-H** GRN derived from the regulons highlighted in **B-D**. (**E**) mouse E18 glycine ACs, (**F**) mouse E18 cones, (**G**) chicken E18 MG, and (**H**) Zebrafish 72hpf HC. The networks show the interplay among genes and regulators within the respective cell classes. Purple nodes represent regulators, gray nodes represent target genes. Red edges indicate regulatory interactions between regulators, gray edges highlight regulatory relationships of regulators with other genes. The layout of the networks is based on centrality, emphasizing the significance of certain nodes within the GRN.

Based on regulon enrichment in specific cell classes, we reconstructed GRN for representative cell class in each stage (Figures 3E-3H, Supplementary Figures S3H–S3Q, S4B– S4G, S5E–S5M). As expected, the regulators within these regulons exhibited pronounced interactions, implying a collaborative effort to preserve cell identity and foster cell differentiation. In glycine ACs in the mouse E18 stage, hub regulators such as Tfap2c and Pou3f1 significantly influenced the GRN (Figure 3E). In cones, Crx, Thrb, Zfp444, and Lcorl (Figure 3F) had multiple regulatory ties. In MG in the chicken E18 stage, SMAD5 and PAX2 were interacting with SOX9 and JUN (Figure 3G). In the zebrafish 72 hpf HC GRN (Figure 3H), regulators such as onecut2, lhx1a, hmx3a, formed a grouped pattern, suggesting that cooperate in certain biological functions.

### Identification of conserved regulators and regulons in mouse, chicken, and zebrafish retina

Given the conservation and comparability of the transcriptional profiles in retinal cells across the three species, we sought to determine if their GRNs were conserved. First, we identified 530 unique cell class-specific regulators from the three species, yielding 250 regulators from mouse, 56 from chicken, and 224 from zebrafish. Then, we examined the overlap of cell class-specific regulators among these species (Supplementary Figure S6A). We identified 10 regulators that were consistently active in all three species, including Gbx2-homolog, Lhx9-homolog, and Pbx1-homolog in ACs and Sox9-homolog in MG. We further confirm their expression specificity across the three species by including human retina organoid development dataset (Figures 4A and 4B, Supplementary Figures S6B-S6D). Specifically, Gbx2 was expressed in a particular subclass of mouse ACs (43), and its homologs were also expressed in ACs across other two species (Figure 4A). However, its human homolog did not appear to be expressed in ACs (Supplementary Figure S6B). Similarly, the Maf-homolog demonstrated were expressed in ACs across all four species (Figure 4B), but its role remains to be explored. We also observed overlapped active regulators among two species in distinct retinal cell classes and further confirmed their expression specificity. For example, HMX3 (hmx3a) and Dmbx1 (dmbx1a), were specifically expressed in chicken and zebrafish HCs and BCs, respectively, but not in human and mouse (Supplementary Figure S7A and S7C). The species-specific expression of HMX3 (hmx3a) likely because of the absence of Type 2 HCs in human and mouse (Supplementary Figure S7B) (48).

**Figure 4.**
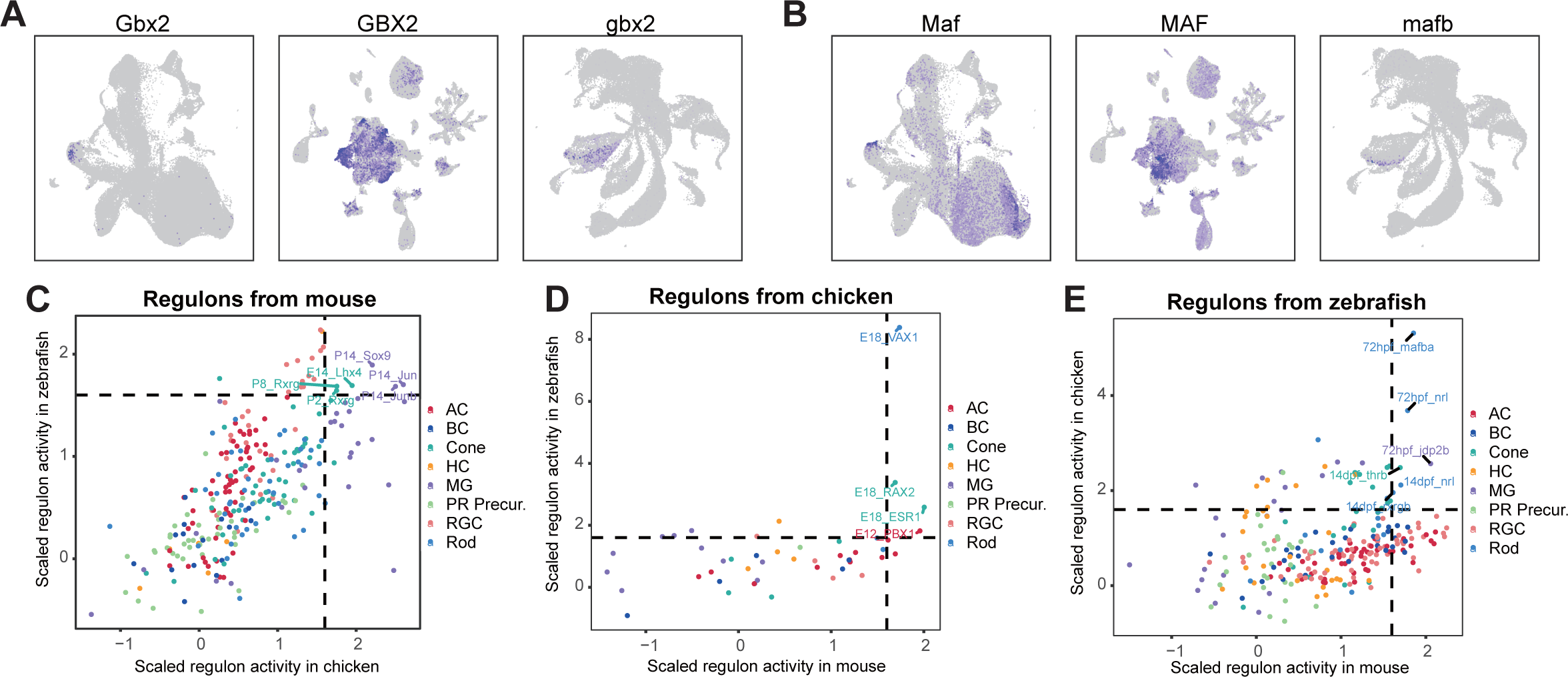
Cross-species analysis of regulators and regulons in homologous retinal cells. **A, B** UMAP expression visualizations of Gbx2 (**A**) and Maf (**B**) among the shared regulators that are specifically active in ACs. Order from left to right: mouse, chicken, zebrafish. Blue indicates high expression, grey indicates low expression. **C-E** Average scaled activity of cell class-specific regulons generated from mouse (**C**), chicken (**D**) and zebrafish (**E**) in the homologous cell class from other two species. Dashed lines at 1.6 (both horizontal and vertical) indicate the thresholds for conserved regulons.

Having closely examined the conservation of regulators across the retinas of the three species, we expanded our analysis to the overarching regulatory networks they orchestrate. We employed AUCell (10) to measure the activity of a cell class-specific regulon from one species across all cell classes in the other two species and computed the average scaled regulon activity for each cell class (see Methods). If a cell class-specific regulon presented a high average scaled activity in its homolog cell class, it was considered a conserved regulon. With an average scaled activity threshold of 1.6, we found 16 conserved regulons: six from mouse (Figure 4C), four from chicken (Figure 4D), and six from zebrafish (Figure 4E). Among the conserved regulons, we found that the Lhx4 regulon (Supplementary Figure S8), which was highly active in mouse cones at E14 stage (Figure 4C), contained several genes co-expressed with well-known marker genes such as *Thrb* -homolog, including *Prom1* -homolog and *Gabrb3* -homolog. Mutation of *GABRB3* has been previously linked to human early infantile epileptic encephalopathy (49). Similarly, we observed that the PBX1 regulon (Supplementary Figure S9), active in chicken ACs at E12 stage (Figure 4D), showed conserved expression of the *Auts2* -homolog across all three species. Mutations in *AUTS2* are strongly associated with autism spectrum disorders (50). The VAX1 regulon (Supplementary Figure S10), active in chicken rods at the E18 stage, included gene ENSGALG00000051404 (si:ch211-113d22.2), which showed rod-specific expression. However, its mouse ortholog was not identified.

### Identification of putative drivers in mouse and zebrafish retinal development

We identified an array of cell class-specific regulators, some of which were conserved across the three species. This led us to question their regulatory roles during developmental processes. To deduce differentiation trajectories for both mouse and zebrafish, we first calculated RNA velocities using scVelo (14) and visualized them through UMAP. The resulting embedding stream plot demonstrated a pronounced motion of RPCs in the direction of mature retinal cell classes, suggesting that RNA velocities capture the critical information on putative drivers of differentiation (Figures 5A and 5B). Additionally, the observed trajectory from BC to rods may be attributed to their similar transcriptional profiles, which could result in overlapping projections in the UMAP space. Based on combined RNA velocities and transcriptional similarity, we determined the fate probability of each cell as well as the putative driver genes guiding their differentiation into specific cell classes (see Methods). We visualized the top 50 correlated putative driver genes for each cell class in the two species (Supplementary Figures S11 and S12). To assess the biological functions of the putative driver genes across mouse and zebrafish, we conducted Gene Ontology (GO) analysis focusing on the biological process (BP) category for the top 100 correlated driver genes in each species independently (see Methods, Supplementary Figure S13). The BPs of driver genes closely aligned with the specific functions of the corresponding cell classes, and homologous cell classes across species shared similar BPs, underscoring the existence of conserved driver genes.

**Figure 5.**
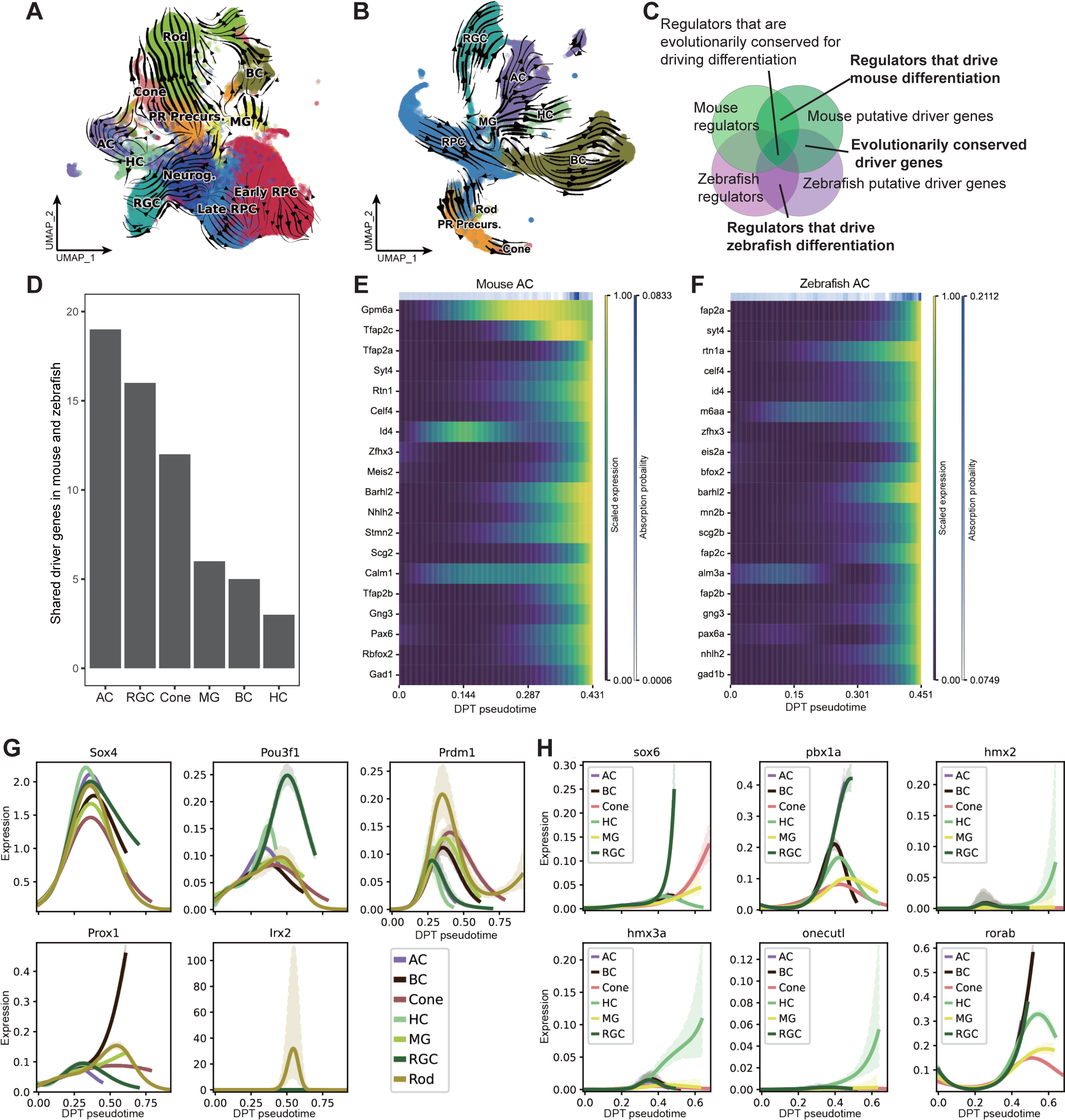
Analysis of RNA velocities, shared putative driver genes, and regulators that drive differentiation in retinal cell development of mouse and zebrafish. **A, B** UMAP-based projections of RNA velocities for retinal cell differentiation in mouse (**A**) and zebrafish (**B**). **C** Venn diagram illustrating the relationships among mouse putative driver genes, mouse regulators, zebrafish regulators, and zebrafish putative driver genes. The intersections highlight (1) shared putative driver genes between mouse and zebrafish and (2) cell class-specific regulators that drive retinal cell development in mouse and zebrafish. **D** Bar chart representing the number of shared putative driver genes among the top 100 cell class-specific driver genes in both mouse and zebrafish. **E, F** Expression heatmaps of shared driver genes in ACs across diffusion pseudotime for mouse (**E**) and zebrafish (**F**). **G, H** Cell class-specific regulators that drive differentiation in mouse (**G**) and zebrafish (**H**).

We further identified two kinds of putative differentiation drivers (Figure 5C): 1) putative differentiation drivers shared between mouse and zebrafish and 2) regulators that drive differentiation in mouse and zebrafish.

We examined the intersection of top 100 putative driver genes in homologous cell classes and observed an overlap, particularly in ACs, RGCs, and cones (Figure 5D). Specifically, 19 shared driver genes were identified in ACs in both species, followed by 16 in RGCs and 12 in cones. Among the shared driver genes, *Pax6* -homolog, *Gng3* -homolog, *Stmn2* -homolog, *Rtn1* -homolog, drove both AC and RGC development (Figures 5E, 5F, and Supplementary Figure S14A, 14B), suggesting that they may be modulated by similar signaling pathways in the early stages of retinal development. *Celf4* -homolog and *Gad1* -homolog were identified as specific drivers of AC differentiation (Figures 5E and 5F), whereas *Elavl4* -homolog, and *Mapt*-homolog were specific drivers of RGC differentiation (Supplementary Figures S14A and S14B). *Guk1* -homolog and *Cuedc1* -homolog were important drivers in cones (Supplementary Figures S14C and S14D). Finally, *Car14* -homolog and *Glul* -homolog were identified as important drivers of MG differentiation (Supplementary Figures S14E and S14F).

To determine cell-class-specific regulators driving retinal cell development, we cross-referenced the top 100 correlated driver genes for each cell class with cell class-specific regulators identified using SCENIC. In mouse, 37 regulators were identified, including widely known ones such as Nrl for rods and Rxrg in cones. We also identified regulators that have not been reported to drive differentiation (Figure 5G). Sox4, Pou3f1, and Prdm1 expression consistently increased across all cell classes during development, with Pou3f1 showing the highest rise in RGCs and Prdm1 the most significant increase in rods. In contrast, Prox1 and Irx2 showed differentiation-dependent increases in expression, with Prox1 increasing exclusively in BCs and Irx2 only in rods. Using a similar approach for zebrafish, 44 regulators were found, including established ones such as klf7a in RGCs and pax6a in ACs. We discovered regulators not previously reported to regulate differentiation (Figure 5H). The expression of sox6 increased in both RGCs and cones during development, whereas pbx1a exhibited a pronounced increase in ACs and RGCs. The expression of hmx2, hmx3a, and onecutl increased only in HCs, and rorab expression showed an increasing trend in ACs, BCs, HCs, and RGCs as development progressed.

Notably, some TFs had already been flagged as cell class-specific regulators in our preceding SCENIC analysis, validating their critical roles in cellular function and identity. For instance, in ACs, TFs such as *Tfap2c* -homolog (Figures 5E and 5F) were not only identified as shared driver genes but also appeared in the SCENIC-identified list of regulators in both mouse and zebrafish. TFs such as *Nhlh2* -homolog were shared driver genes but not listed as regulators in the SCENIC analysis (Figures 5E and 5F).

### Identification of conserved expression patterns between vertebrate retina and *Ciona* PR-related cells

To expand our evolutionary understanding beyond the conserved regulatory genes identified in the mouse, chicken, and zebrafish retinas, we included an invertebrate *Ciona* in our comparative study. We first identified a set of conserved DEGs in the retinal cells of vertebrates. The top 10 conserved DEGs are consistently expressed in the corresponding homologous cells (Figure 6A). Then, we collected scRNA-seq data from *C. intestinalis*, extracted eight PR-related cell types (Figure 6B), and identified DEGs in these cell types. We analyzed the intersection of DEGs between vertebrate retinal cell classes and *Ciona* PR-related cells based on the 5894 orthogroups generated using Orthofinder (Figure 6C, see Methods) (38). Specifically, if DEGs in a certain type of *Ciona* PR-related cells have orthologous DEGs in a certain class of vertebrate retinal cell, these two cell types may have a homologous relationship. Twenty-one orthologs of the 23 DEGs in vertebrate retinal cells also displayed differential expression in PR-related cells in *Ciona* (Supplementary Table S7). Specifically, in the Rx+ aSV and Ci-VP+ pSV_cells of *Ciona*, four and six orthologs were differentially expressed in vertebrate PRs and RGCs, respectively (Figure 6D). A previous study showed that RX is specifically expressed in developing PRs of Ciona, indicating that the Rx+ aSV cells correspond to developing PRs in *Ciona* (51).

**Figure 6.**
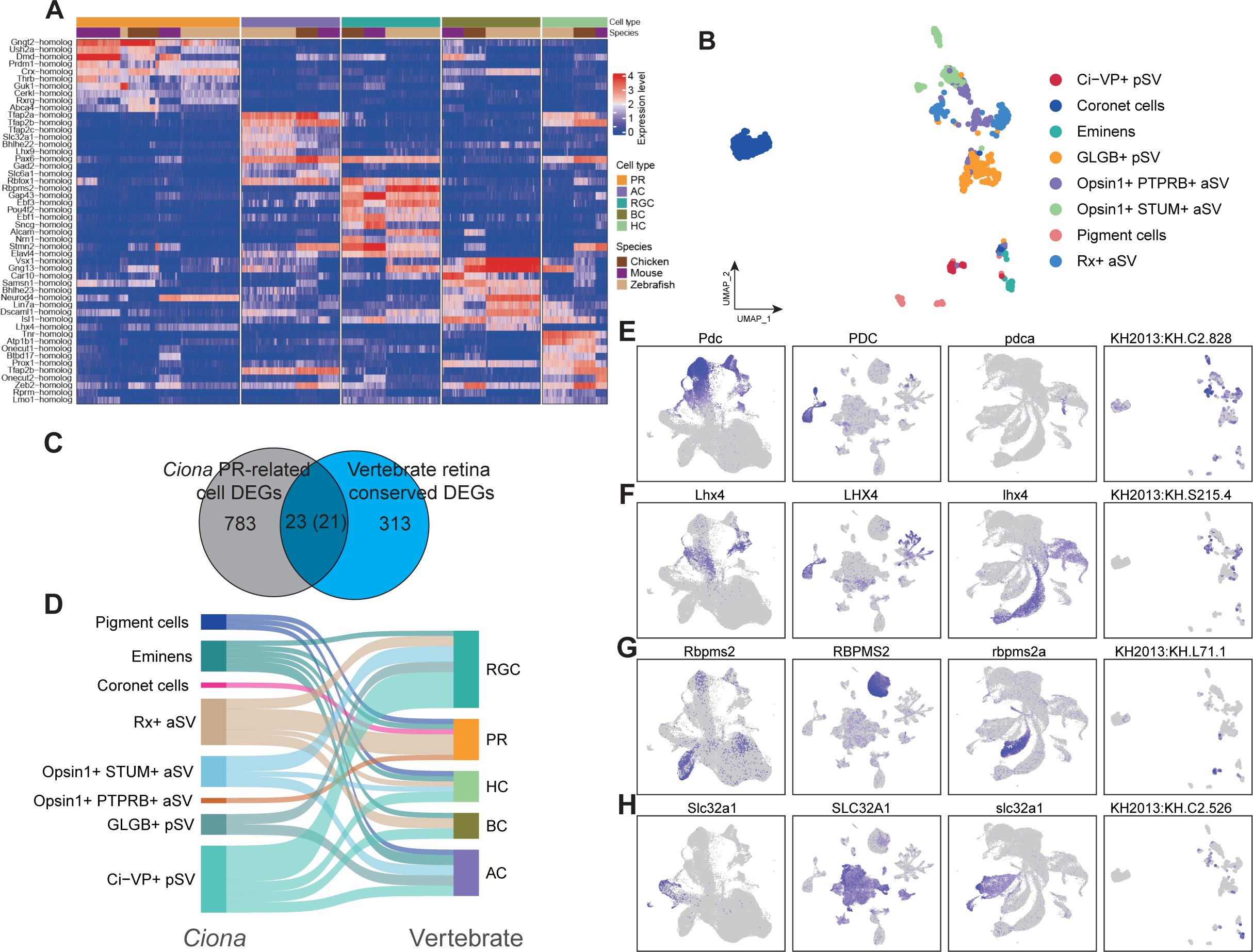
Exploration of the origin of the vertebrate retina. **A** Heatmap of the top 10 conserved DEGs in homologous cell classes of the vertebrate retina. Red indicates high expression, blue indicates low expression. **B** UMAP visualization of scRNA-seq data from *Ciona* PR-related cells. **C** Venn diagram showing overlapping DEGs between vertebrate retina cells and Ciona PR-related cells, used to infer the origin of vertebrate retina. **D** Sankey diagram showing shared DEGs between *Ciona* PR-related cells and vertebrate retinal cells. **E-H** UMAP visualizations of gene expression levels across the four species. Each panel represents the expression of a specific gene: (**E**) *Pdc* -orth, (**F**) *Lhx4* - orth, (**G**) *Rbpms2* -orth, and (**H**) *Slc32a1* -orth. Order from left to right: mouse, chicken, zebrafish, and *Ciona*. Blue indicates high expression, gray indicates low expression.

We further investigated the functions of these overlapped orthologs. Rx+ aSV cells specifically express KH.C2.828 and KH.C1.346, orthologs to vertebrate PR-specific genes Phosducin (PDC) and Prom1, respectively (Figure 6E, Supplementary Figures S15A). PDC modulates G-protein-coupled receptor signaling (52), and Prom1 mutations link to retinal degeneration (53). This observation suggests that *Ciona* PRs shared similar light transduction mechanisms with vertebrate PRs. Additionally, Rx+ aSV cells also specifically expressed KH.S215.4 and KH.L152.2, which correspond to the transcription factors Lhx4 and Isl2, key regulators of vertebrate BCs, cones, and RGCs (Figure 6F, Supplementary Figures S15B) (54, 55). This finding indicates that *Ciona* PRs share a regulatory program similar to that of vertebrate PRs. Furthermore, Ci-VP+ pSV cells expressed KH.L71.1 and KH.C14.291, orthologs to Rbpms2 and Dach2, which were DEGs in RGCs and HCs (Figure 6G, Supplementary Figures S15C). KH.L24.10 and KH.C12.301, orthologs to Ebf1 and Fstl4, were expressed in Opsin1+ STUM+ aSV and Ci-VP+ pSV cells (Supplementary Figures S15D and S15E). Rbpms2 and Fstl4 are markers for RGCs (56, 57), while Ebf1 regulates RGC differentiation (58). These findings suggest that Ci-VP+ pSV cells shares functional similarities with vertebrate RGCs. Finally, three DEGs from *Ciona* inhibitory neuron eminens had orthologs specifically expressed in various vertebrate retina cell classes. KH.C2.526 and KH.L50.5, orthologs to Slc32a1 (VGAT) and Slc24a2 (NCKX2), were expressed in vertebrate ACs and cones, respectively (Figure 6H, Supplementary S15F). Slc32a1 loads inhibitory amino acids into vesicles (59), while Slc24a2 regulates ion levels for light response (60). KH.S761.6, orthologous to Gad2, is a marker for GABAergic neurons (61) and was expressed in vertebrate ACs and HCs (Supplementary S15G).

We further confirmed the expression patterns of these overlapped orthologs in their human retina development organoid dataset. As expected, these conserved genes were also specifically expressed in the corresponding cell class (Supplementary Figures S16A-S16E), demonstrating their unique function in retinal cells.

## DISCUSSION

In this study, we comprehensively explored the transcriptional landscape of vertebrate retinal cells across three representative species, unveiling transcriptomic conservation suggestive of shared regulatory mechanisms. Our detailed investigation into cell class-specific regulatory networks provided insights into the inherent regulatory dynamics of retinal cells, emphasizing the role of conserved regulators and regulons in evolutionary terms. Furthermore, comparative analysis with the invertebrate *Ciona* underscored potential evolutionary relations, hinting at shared ancestral origins of certain retinal cells.

Our findings corroborate the idea that the transcriptomes of homologous cells are more similar across species than they are to other cell classes within the same species. This observation is consistent with a recent study surveying birds, mammals, reptiles, fish, and lamprey (9). This not only affirms evolutionary conservation but also suggests the pivotal role of these cells in visual function across diverse vertebrates. While our study provided insights into general transcriptomic similarities among homologous cells in vertebrates, we did not delve into the finer subclasses of these cells. For instance, 40 types of RGCs have been identified in mouse (57), and more than 30 types in zebrafish (62). Future studies based on more refined cell-type classifications may unveil additional layers of transcriptomic and functional complexity, providing new avenues for the exploration of retinal biology and evolution.

Our application of SCENIC uncovered 690 high-quality regulons and their corresponding 530 regulators. The smaller number of regulators identified in chicken can be attributed to the fewer time points analyzed for this species. These regulators showed a high overlap with regulators previously reported in the literature (Supplementary Tables S4-S6). For example, previous research has established the importance of Crx, Otx2, and Thrb in cone specification (63). Our study extends this by demonstrating their high interaction with Zfp444 and Lcorl in cones at the mouse E18 stage, suggesting potential roles for these genes in cone specification (Figure 3F). Additionally, while lhx1a is known as an HC marker gene(48), and onecut2 is a regulator of HC fate (64), we discovered their interaction with hmx3a and rxraa in zebrafish HCs at 72 hpf stage (Figure 3H). The functions of hmx3a and rxraa may also be related to the identity and specification of zebrafish HCs. In summary, this extensive repository of regulons and regulators, many of which remain to be explored, represents a rich resource for future investigations into the intricate mechanisms of vertebrate retinal biology.

We pinpointed 10 conserved regulators and 16 conserved regulons in homologous cells across vertebrates. The relatively modest number of overlapping regulators and regulons can be attributed to several factors: the inherent sparsity of single-cell RNA-seq data, the exclusion of some transcription factors when constructing the ranking database (see Methods), the limited number of chicken datasets, and the stringent criteria applied to filter high quality regulons. The conservation of these regulators suggests their fundamental importance to basic cellular functions, while also hinting at unexplored, potentially critical roles. For example, Gbx2, although proven to be expressed in a subtype of GABAergic ACs (43), has yet to have its function established in the AC of chicken and zebrafish. The conserved regulons we identified not only enhance our understanding of the fundamental functions of the retina but offers insights into complex brain functions. For instance, we observed a conserved expression of Gabrb3 and Auts2 in cones and ACs across the three species from the conserved regulons (Supplementary Figures S8 and S9). While these genes has been linked to human brain diseases like human early infantile epileptic encephalopathy and autism spectrum disorders (49, 50), their specific functions in these diseases remain to be further investigated. Studying the functions of these conserved genes in a simpler model, such as the zebrafish retina, provides a more feasible approach than exploring them in the complex environment of the human brain.

Our RNA velocity analysis in mouse and zebrafish significantly advanced our comprehension of genes driving retinal differentiation in vertebrates. The putative driver genes we found in mouse were similar to those identified in a previous study (6), which used pseudo-time trajectory inference to predict differentiation genes. For example, both our study and the previous study identified important genes such as Pax6 and Sox4 in ACs. Additionally, we identified new putative driver genes such as Nhlh2, providing new perspectives on the differentiation of the mouse retina. In addition, we identified conserved putative driver genes between mouse and zebrafish, overlapping with previously identified marker genes, and demonstrated their intricate involvement in establishing cell identity. For example, Gad1, a known marker for ACs (65), was predominantly expressed in the terminal stages of differentiation (Figure 5E and 5F); similar patterns were also observed in other putative driver genes such as Celf4. This not only corroborates the role of Gad1 as a marker gene but highlights its importance across species in establishing cell identity. In addition, we uncovered a set of regulators that drive differentiation, which allowed us to pinpoint transcription factors and regulatory networks pivotal in guiding cellular differentiation pathways. A recent study reported the regulatory role of Pou3f1 in RGC differentiation (66). This finding closely aligns with our results (Figure 5G), supporting the validity of our identified regulators.

Our comparative analysis with the invertebrate species *Ciona* allowed us to identify DEGs shared between vertebrate retinal cells and *Ciona* PR-related cells. These shared genes have been widely used as markers for vertebrate retinal cells though their functions are not yet fully understood. For instance, *Rbpms2* is an RNA binding protein with multiple splicing. It has been used as a marker for vertebrate RGCs, but its specific role in these cells is still unknown (56). Considering the compact genome of *Ciona* (37), exploring the regulatory programs of their *Ciona* orthologs could yield valuable insights into their functional conservation and evolutionary dynamics. Furthermore, through these shared DEGs, we identified corresponding cell types between vertebrate retina and *Ciona* PR-related cells, such as vertebrate PRs and *Ciona* Rx+ aSV cells, as well as vertebrate RGCs and *Ciona* Ci-VP+ cells. This correlation helps us better investigate the simple patterns of information transmission within retina, offering insights into more complex visual systems. However, it is important to note that our conclusions about ancestral relationships are drawn from a small number of conservative genes. Expanding this analysis to include a broader range of genes and other types of molecular data may provide a more comprehensive view of evolutionary relationships and might yield additional insights into the ancestral forms of these retinal cells.

## DATA AVAILABILITY

FASTQ data from scRNA-seq retinal cells were retrieved from the GEO database for the four species: mouse, chicken, and zebrafish, *Ciona* (GSE118614, GSE159107, GSE122680, GSE131155). Codes generated for this project is available on GitHub (https://github.com/xzengComBio/VertebrateRetina_scRNAseq).

## SUPPLEMENTARY DATA

Supplementary Data are available at NAR online.

## AUTHOR CONTRIBUTIONS

Conceptualization, X.Z., F.G., T.G.K., and K.N.; Data Curation, X.Z.; Formal Analysis, X.Z.; Investigation, X.Z., F.G., C.Y., and T.G.K.; Writing – Original Draft, X.Z. and M.L.; Writing – Review & Editing, X.Z., F.G., Y.C., M.L., T.G.K., and K.N.; Funding Acquisition, T.G.K. and K.N.; Resources, K.N.; Supervision, T.G.K. and K.N.

## ACKNOWLEDGEMENTS

Computational resources were provided by the supercomputer system SHIROKANE at the Human Genome Center, Institute of Medical Science, the University of Tokyo.

