## Supplemental Figures for "Comparative single-cell transcriptomic analysis reveals putative differentiation drivers and potential origin of vertebrate retina"

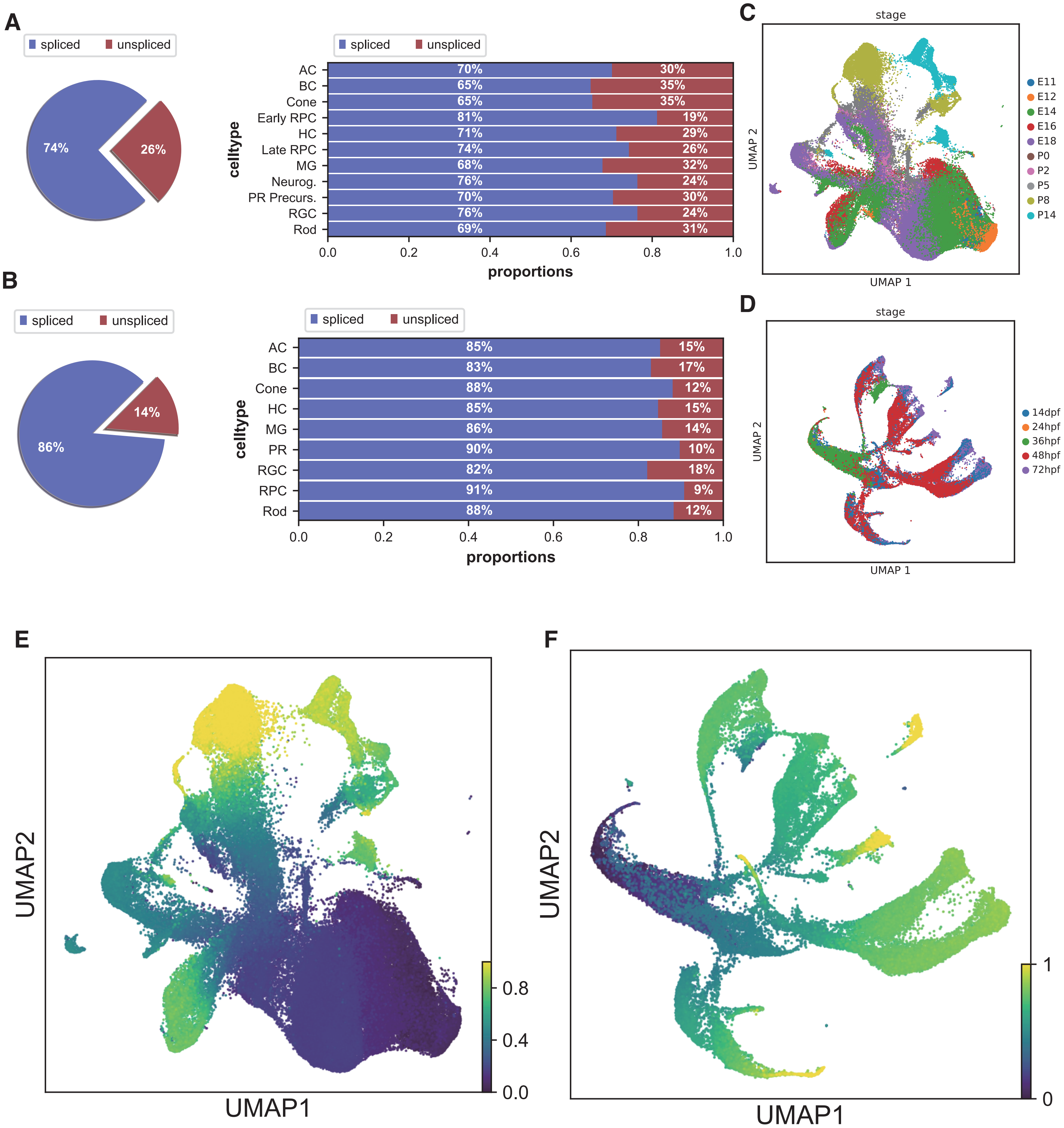


**Figure S1 RNA velocity preprocessing in mouse and zebrafish retinal cells.**
**A, B** Analysis of unspliced and spliced RNA proportions. (Left) Pie chart representing overall proportions of unspliced vs. spliced RNA. (Right) Barplot detailing unsplice/splice RNA proportions for each cell type. (**A**) mouse (**B**) zebrafish. **C, D**: UMAP visualizations of cells colored by stage. (**C**) mouse, (**D**) zebrafish. **E, F**:UMAP visualizations based on inferred DPT (diffusion pseudotime). (**E**) mouse (**F**) zebrafish.

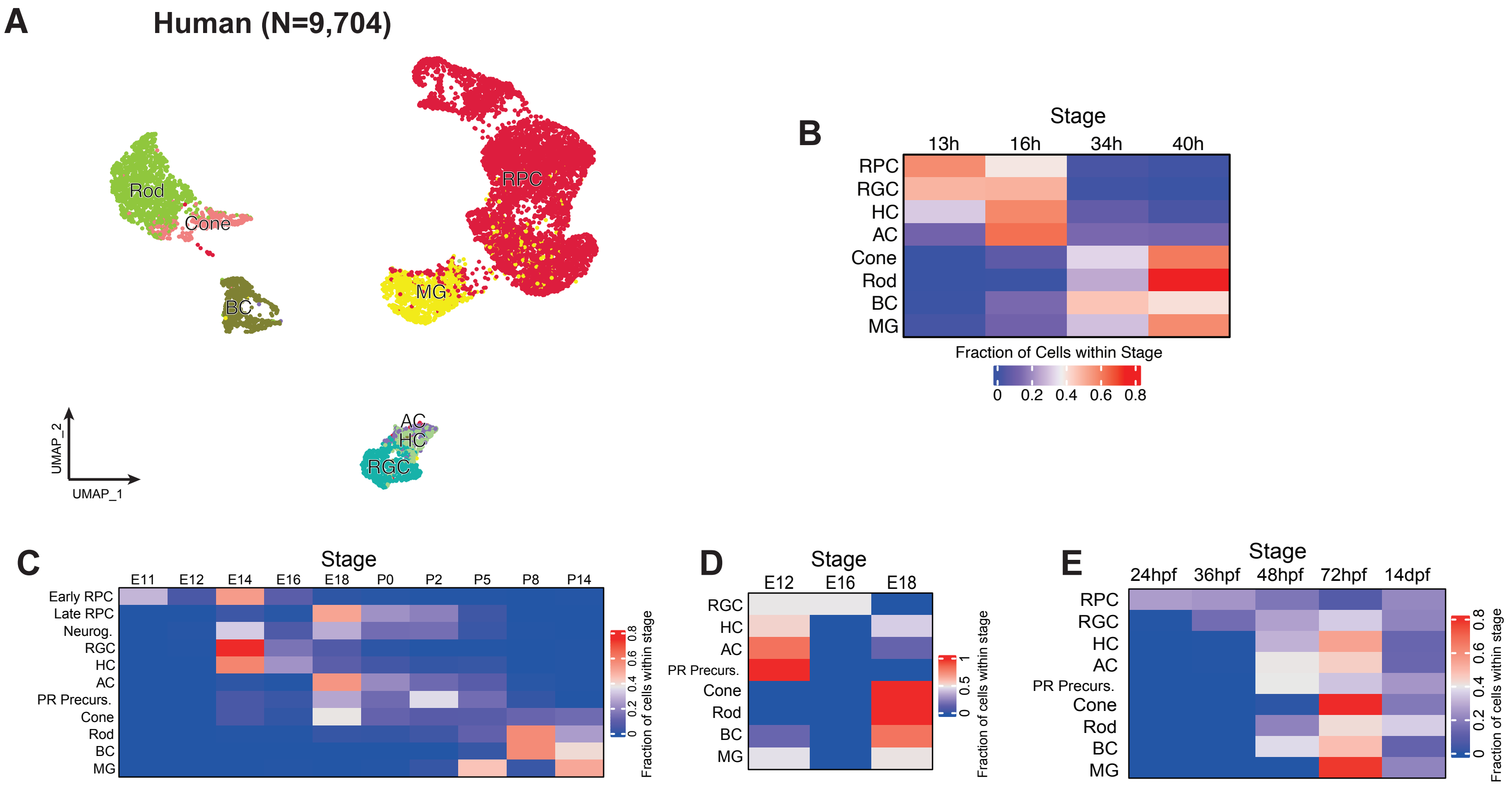


**Figure S2 Cell type distributions across developmental stages in human retinal organoids and retinas from other species**

**A**: UMAP plots of retinal scRNA-seq data from human retina organoid development. **B-E**: Proportions of cell types within each stage for human (**B**), mouse (**C**), chicken (**D**), and zebrafish (**E**). Red indicates a high proportion, blue indicates a low proportion.

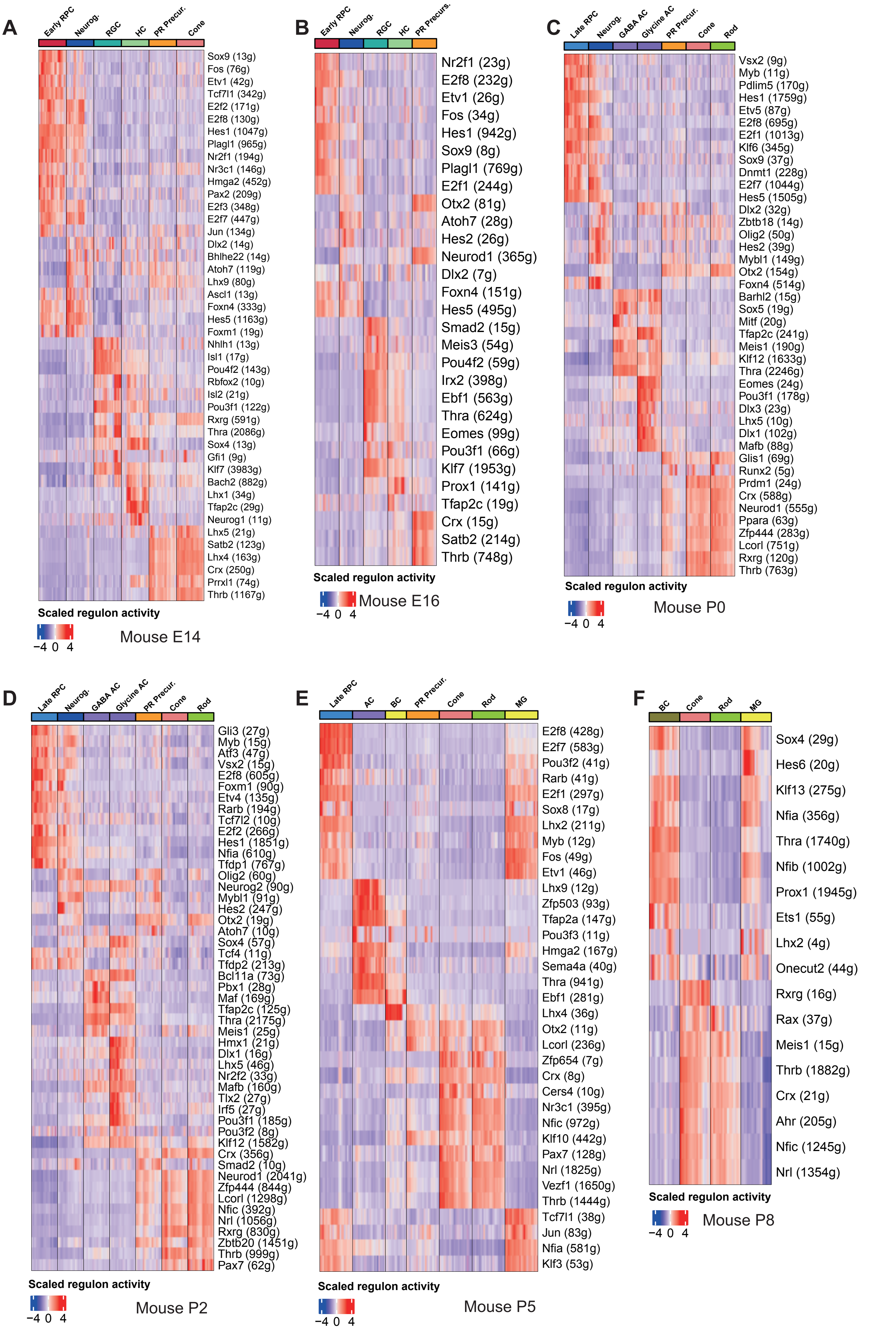
A
Early RPC
Neurog.
RGC
HC
PR Precur.
Cone
Sox9 (13g)
Fos (76g)
Etv1 (42g)
Tcf7l1 (342g)
E2f2 (171g)
E2f8 (130g)
Hes1 (1047g)
Plagl1 (965g)
Nr2f1 (194g)
Nr3c1 (146g)
Hmga2 (452g)
Pax2 (209g)
E2f3 (348g)
E2f7 (447g)
Jun (134g)
Dlx2 (14g)
Bhlhe22 (14g)
Atoh7 (119g)
Lhx9 (80g)
Ascl1 (13g)
Foxn4 (333g)
Hes5 (1163g)
Foxm1 (19g)
Nhlh1 (13g)
Isl1 (17g)
Pou4f2 (143g)
Rbfox2 (10g)
Isl2 (21g)
Pou3f1 (122g)
Rxrg (591g)
Thra (2086g)
Sox4 (13g)
Gfi1 (9g)
Klf7 (3983g)
Bach2 (882g)
Lhx1 (34g)
Tfap2c (29g)
Neurog1 (11g)
Lhx5 (21g)
Satb2 (123g)
Lhx4 (163g)
Crx (250g)
Prrxl1 (74g)
Thrb (1167g)
Scaled regulon activity
−4 0 4
Mouse E14
B
Early RPC
Neurog.
RGC
HC
PR Precurs.
Nr2f1 (23g)
E2f8 (232g)
Etv1 (26g)
Fos (34g)
Hes1 (942g)
Sox9 (8g)
Plagl1 (769g)
E2f1 (244g)
Otx2 (81g)
Atoh7 (28g)
Hes2 (26g)
Neurod1 (365g)
Dlx2 (7g)
Foxn4 (151g)
Hes5 (495g)
Smad2 (15g)
Meis3 (54g)
Pou4f2 (59g)
Irx2 (398g)
Ebf1 (563g)
Thra (624g)
Eomes (99g)
Pou3f1 (66g)
Klf7 (1953g)
Prox1 (141g)
Tfap2c (19g)
Crx (15g)
Satb2 (214g)
Thrb (748g)
Scaled regulon activity
−4 0 4
Mouse E16
C
Late RPC
Neurog.
GABA AC
Glycine AC
PR Precur.
Cone
Rod
Vsx2 (9g)
Myb (11g)
Pdlim5 (170g)
Hes1 (1759g)
Etv5 (87g)
E2f8 (695g)
E2f1 (1013g)
Klf6 (345g)
Sox9 (37g)
Dnmt1 (228g)
E2f7 (1044g)
Hes5 (1505g)
Dlx2 (32g)
Zbtb18 (14g)
Olig2 (50g)
Hes2 (39g)
Mybl1 (149g)
Otx2 (154g)
Foxn4 (514g)
Barhl2 (15g)
Sox5 (19g)
Mitf (20g)
Tfap2c (241g)
Meis1 (190g)
Klf12 (1633g)
Thra (2246g)
Eomes (24g)
Pou3f1 (178g)
Dlx3 (23g)
Lhx5 (10g)
Dlx1 (102g)
Mafb (88g)
Glis1 (69g)
Runx2 (5g)
Prdm1 (24g)
Crx (588g)
Neurod1 (555g)
Ppara (63g)
Zfp444 (283g)
Lcorl (751g)
Rxrg (120g)
Thrb (763g)
Scaled regulon activity
−4 0 4
Mouse P0
D
Late RPC
Neurog.
GABA AC
Glycine AC
PR Precur.
Cone
Rod
Gli3 (27g)
Myb (15g)
Atf3 (47g)
Vsx2 (15g)
E2f8 (605g)
Foxm1 (90g)
Etv4 (135g)
Rarb (194g)
Tcf7l2 (10g)
E2f2 (266g)
Hes1 (1851g)
Nfia (610g)
Tfdp1 (767g)
Olig2 (60g)
Neurog2 (90g)
Mybl1 (91g)
Hes2 (247g)
Otx2 (19g)
Atoh7 (10g)
Sox4 (57g)
Tcf4 (11g)
Tfdp2 (213g)
Bcl11a (73g)
Pbx1 (28g)
Maf (169g)
Tfap2c (125g)
Thra (2175g)
Meis1 (25g)
Hmx1 (21g)
Dlx1 (16g)
Lhx5 (46g)
Nr2f2 (33g)
Mafb (160g)
Tlx2 (27g)
Irf5 (27g)
Pou3f1 (185g)
Pou3f2 (8g)
Klf12 (1582g)
Crx (356g)
Smad2 (10g)
Neurod1 (2041g)
Zfp444 (844g)
Lcorl (1298g)
Nfic (392g)
Nrl (1056g)
Rxrg (830g)
Zbtb20 (1451g)
Thrb (999g)
Pax7 (62g)
Scaled regulon activity
−4 0 4
Mouse P2
E
Late RPC
AC
BC
PR Precur.
Cone
Rod
MG
E2f8 (428g)
E2f7 (583g)
Pou3f2 (41g)
Rarb (41g)
E2f1 (297g)
Sox8 (17g)
Lhx2 (211g)
Myb (12g)
Fos (49g)
Etv1 (46g)
Lhx9 (12g)
Zfp503 (93g)
Tfap2a (147g)
Pou3f3 (11g)
Hmga2 (167g)
Sema4a (40g)
Thra (941g)
Ebf1 (281g)
Lhx4 (36g)
Otx2 (11g)
Lcorl (236g)
Zfp654 (7g)
Crx (8g)
Cers4 (10g)
Nr3c1 (395g)
Nfic (972g)
Klf10 (442g)
Pax7 (128g)
Nrl (1825g)
Vezf1 (1650g)
Thrb (1444g)
Tcf7l1 (38g)
Jun (83g)
Nfia (581g)
Klf3 (53g)
Scaled regulon activity
−4 0 4
Mouse P5
F
BC
Cone
Rod
MG
Sox4 (29g)
Hes6 (20g)
Klf13 (275g)
Nfia (356g)
Thra (1740g)
Nfib (1002g)
Prox1 (1945g)
Ets1 (55g)
Lhx2 (4g)
Onecut2 (44g)
Rxrg (16g)
Rax (37g)
Meis1 (15g)
Thrb (1882g)
Crx (21g)
Ahr (205g)
Nfic (1245g)
Nrl (1354g)
Scaled regulon activity
−4 0 4
Mouse P8

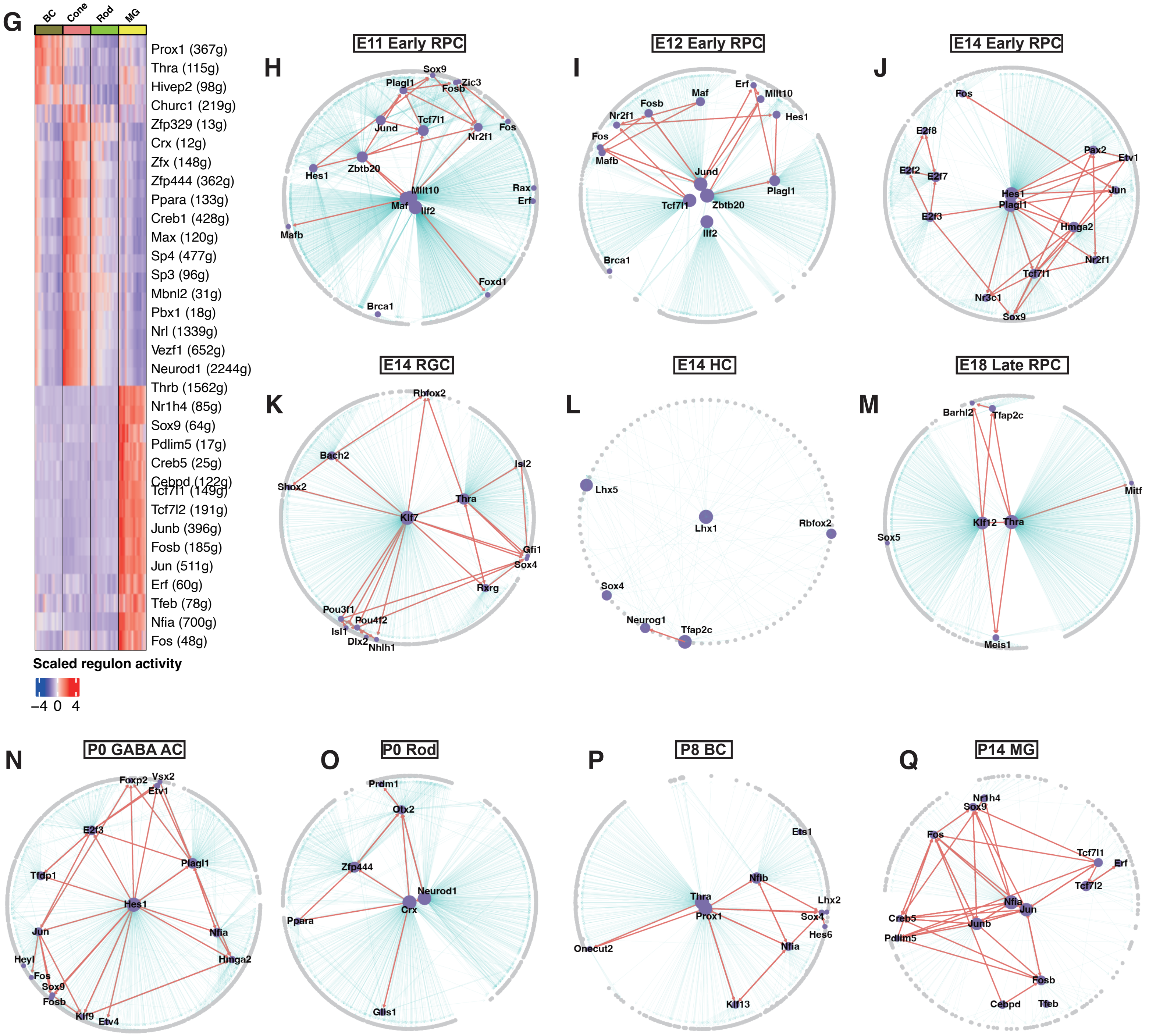


**Figure S3 Reconstruction of GRN in mouse retinal cells at various stages**

**A-G**: Heatmaps of scaled regulon activity across different stages in mouse. (**A**) E14, (**B**) E16, (**C**) P0, (**D**) P2, (**E**) P5, (**F**) P8, (**G**) P14. Each row represents a distinct regulon, while each column denotes a specific cell type. Each regulon is named following the format "TF (n)", where "TF" represents the regulator and "n" denotes the number of target genes in that regulon. Intensity of red signifies high activity, whereas blue indicates low activity. **H-Q:** GRN derived from the regulons in **A-G** and Figure **3B**. (**H**) E11 Early RPC. (**I**) E12 Early RPC. (**J**) E14 Early RPC. (**K**) E14 RGC. (**L**) E14 HC, (**M**) E18 Late RPC, (**N**) P0 GABA AC, (**O**) P0 Rod, (**P**) P8 BC, (**Q**) P14 MG. These networks showcase the interplay among genes and regulators within the respective cell types. Purple nodes represent regulators, while gray nodes indicate target genes. Red edges depict regulatory interactions between regulators, and gray edges highlight regulatory relationships of regulators to other genes.

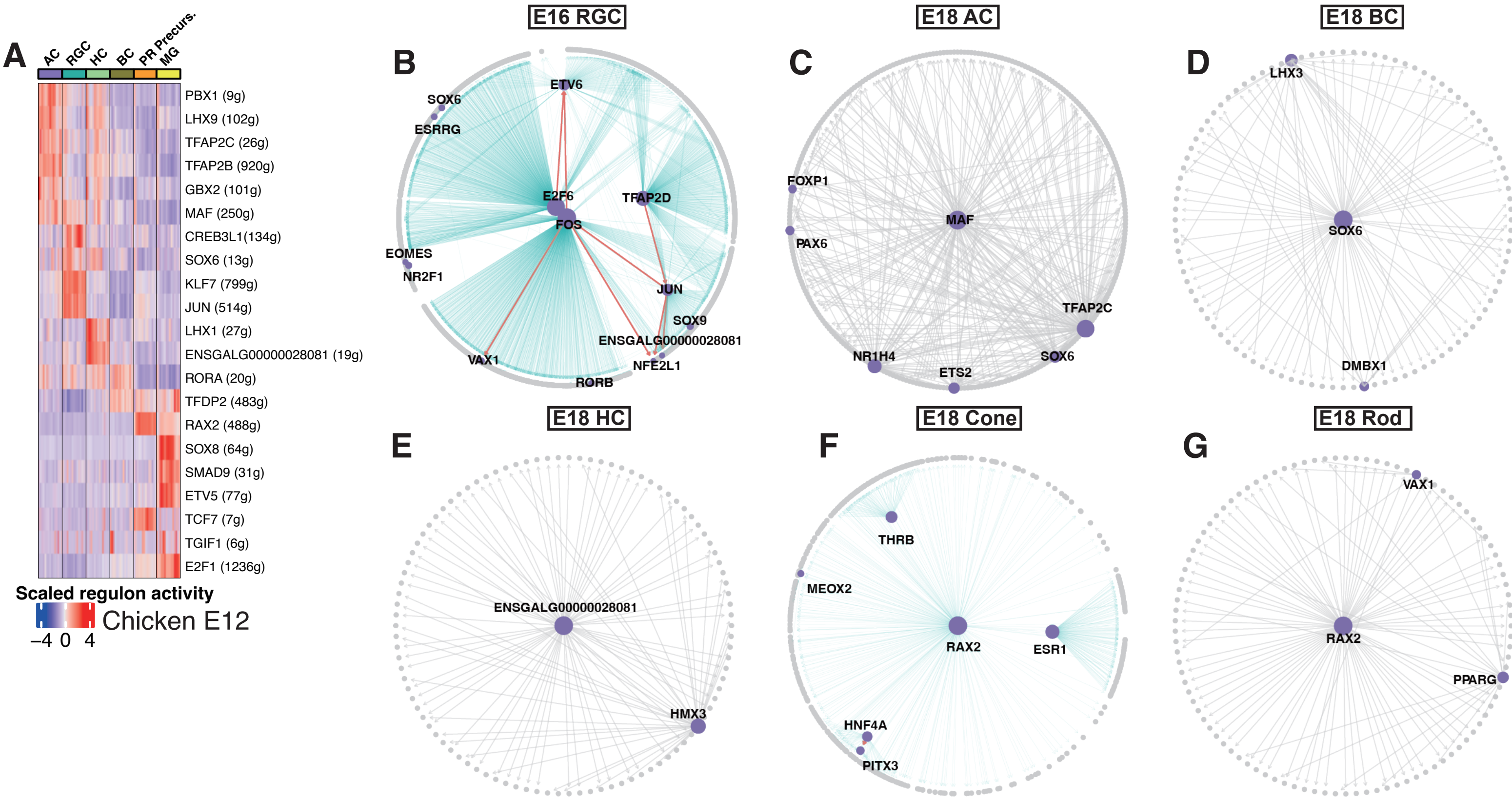


**Figure S4 Reconstruction of GRN in chicken retinal cells**

**A** Heatmaps of scaled regulon activity across in E12. Each row represents a distinct regulon, while each column denotes a specific cell type. Each regulon is named following the format "TF (n)", where "TF" represents the regulator and "n" denotes the number of target genes in that regulon. Intensity of red signifies high activity, whereas blue indicates low activity. **B–G:** GRN derived from the regulons in **A** and Figure **3C**. (**B**) E16 RGC, (**C**) E18 AC, (**D**) E18 BC, (**E**) E18 HC, (**F**) E18 Cone, (**G**) E18 Rod. These networks showcase the interplay among genes and regulators within the respective cell types. Purple nodes represent regulators, while gray nodes indicate target genes. Red edges depict regulatory interactions between regulators, and gray edges highlight regulatory relationships of regulators to other genes.

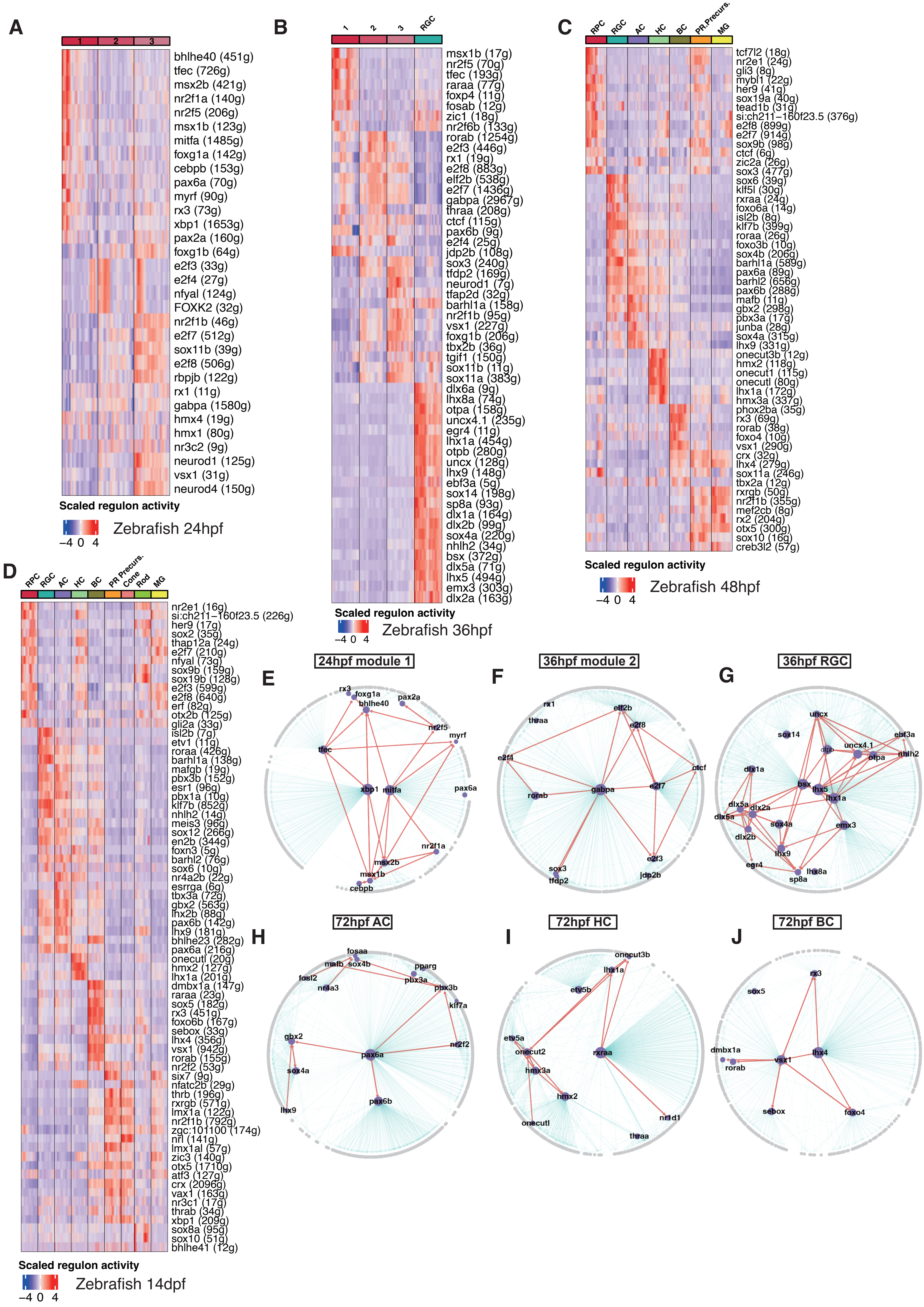
A
1 2 3
bhlhe40 (451g)
tfec (726g)
msx2b (421g)
nr2f1a (140g)
nr2f5 (206g)
msx1b (123g)
mitfa (1485g)
foxg1a (142g)
cebpb (153g)
pax6a (70g)
myrf (90g)
rx3 (73g)
xbp1 (1653g)
pax2a (160g)
foxg1b (64g)
e2f3 (33g)
e2f4 (27g)
nfyal (124g)
FOXK2 (32g)
nr2f1b (46g)
e2f7 (512g)
sox11b (39g)
e2f8 (506g)
rbpjb (122g)
rx1 (11g)
gabpa (1580g)
hmx4 (19g)
hmx1 (80g)
nr3c2 (9g)
neurod1 (125g)
vsx1 (31g)
neurod4 (150g)
Scaled regulon activity
−4 0 4
Zebrafish 24hpf
B
1 2 3 RGC
msx1b (17g)
nr2f5 (70g)
tfec (193g)
raraa (77g)
foxp4 (11g)
fosab (12g)
zic1 (18g)
nr2f6b (133g)
rorab (1254g)
e2f3 (446g)
rx1 (19g)
e2f8 (883g)
elf2b (538g)
e2f7 (1436g)
gabpa (2967g)
thraa (208g)
ctcf (115g)
pax6b (9g)
e2f4 (25g)
jdp2b (108g)
sox3 (240g)
tfdp2 (169g)
neurod1 (7g)
tfap2d (32g)
barhl1a (158g)
nr2f1b (95g)
vsx1 (227g)
foxg1b (206g)
tbx2b (36g)
tgif1 (150g)
sox11b (11g)
sox11a (383g)
dlx6a (9g)
lhx8a (74g)
otpa (158g)
uncx4.1 (235g)
egr4 (11g)
lhx1a (454g)
otpb (280g)
uncx (128g)
lhx9 (148g)
ebf3a (5g)
sox14 (198g)
sp8a (93g)
dlx1a (164g)
dlx2b (99g)
sox4a (220g)
nhlh2 (34g)
bsx (372g)
dlx5a (71g)
lhx5 (494g)
emx3 (303g)
dlx2a (163g)
Scaled regulon activity
−4 0 4
Zebrafish 36hpf
C
RPC RGC AC HC BC PR.Precurs. MG
tcf7l2 (18g)
nr2e1 (24g)
gli3 (8g)
mybl1 (22g)
her9 (41g)
sox19a (40g)
tead1b (31g)
si:ch211−160f23.5 (376g)
e2f8 (899g)
e2f7 (914g)
sox9b (98g)
ctcf (6g)
zic2a (26g)
sox3 (477g)
sox6 (39g)
klf5l (30g)
rxraa (24g)
foxo6a (14g)
isl2b (8g)
klf7b (399g)
roraa (26g)
foxo3b (10g)
sox4b (206g)
barhl1a (589g)
pax6a (89g)
barhl2 (656g)
pax6b (288g)
mafb (11g)
gbx2 (298g)
pbx3a (17g)
junba (28g)
sox4a (315g)
lhx9 (331g)
onecut3b (12g)
hmx2 (118g)
onecut1 (115g)
onecutl (80g)
lhx1a (172g)
hmx3a (337g)
phox2ba (35g)
rx3 (69g)
rorab (38g)
foxo4 (10g)
vsx1 (290g)
crx (32g)
lhx4 (279g)
sox11a (246g)
tbx2a (12g)
rxrgb (50g)
nr2f1b (355g)
mef2cb (8g)
rx2 (204g)
otx5 (300g)
sox10 (16g)
creb3l2 (57g)
Scaled regulon activity
−4 0 4
Zebrafish 48hpf
D
RPC RGC AC HC BC PR Precurs. Cone Rod MG
nr2e1 (16g)
si:ch211−160f23.5 (226g)
her9 (17g)
sox2 (35g)
thap12a (24g)
e2f7 (210g)
nfyal (73g)
sox9b (159g)
sox19b (128g)
e2f3 (599g)
e2f8 (640g)
erf (82g)
otx2b (125g)
gli2a (33g)
isl2b (7g)
etv1 (11g)
roraa (426g)
barhl1a (138g)
mafgb (19g)
pbx3b (152g)
esrl (96g)
pbx1a (10g)
klf7b (852g)
nhlh2 (14g)
meis3 (96g)
sox12 (266g)
en2b (344g)
foxn3 (5g)
barhl2 (76g)
sox6 (10g)
nr4a2b (22g)
esrrga (6g)
tbx3a (72g)
gbx2 (563g)
lhx2b (88g)
pax6b (142g)
lhx9 (181g)
bhlhe23 (282g)
pax6a (216g)
onecutl (20g)
hmx2 (127g)
lhx1a (201g)
dmbx1a (147g)
raraa (23g)
sox5 (182g)
rx3 (451g)
foxo6b (167g)
sebox (33g)
lhx4 (356g)
vsx1 (942g)
rorab (155g)
nr2f2 (53g)
six7 (9g)
nfatc2b (29g)
thrb (196g)
rxrgb (571g)
lmx1a (122g)
nr2f1b (792g)
zgc:101100 (174g)
nrl (141g)
lmx1al (57g)
zic3 (140g)
otx5 (1710g)
atf3 (127g)
crx (2096g)
vax1 (163g)
nr3c1 (17g)
thrab (34g)
xbp1 (209g)
sox8a (95g)
sox10 (51g)
bhlhe41 (12g)
Scaled regulon activity
−4 0 4
Zebrafish 14dpf
E
24hpf module 1
rx3
foxg1a
bhlhe40
pax2a
nr2f5
myrf
tfec
xbp1
mitfa
pax6a
nr2f1a
msx2b
msx1b
cebpb
F
36hpf module 2
rx1
thraa
elf2b
e2f8
e2f4
ctcf
rorab
gabpa
e2f7
e2f3
sox3
tfdp2
jdp2b
G
36hpf RGC
uncx
sox14
ebf3a
uncx4.1
otpb
otpa
nhlh2
dlx1a
bsx
lhx5
lhx1a
dlx5a
dlx2a
dlx6a
sox4a
emx3
dlx2b
lhx9
egr4
lhx8a
sp8a
H
72hpf AC
fosaa
mafb
sox4b
pparg
fosl2
nr4a3
pbx3a
pbx3b
klf7a
gbx2
nr2f2
pax6a
sox4a
pax6b
lhx9
I
72hpf HC
onecut3b
lhx1a
etv5b
etv5a
onecut2
rxraa
hmx3a
hmx2
nr1d1
onecutl
thraa
J
72hpf BC
rx3
sox5
dmbx1a
rorab
vsx1
lhx4
sebox
foxo4

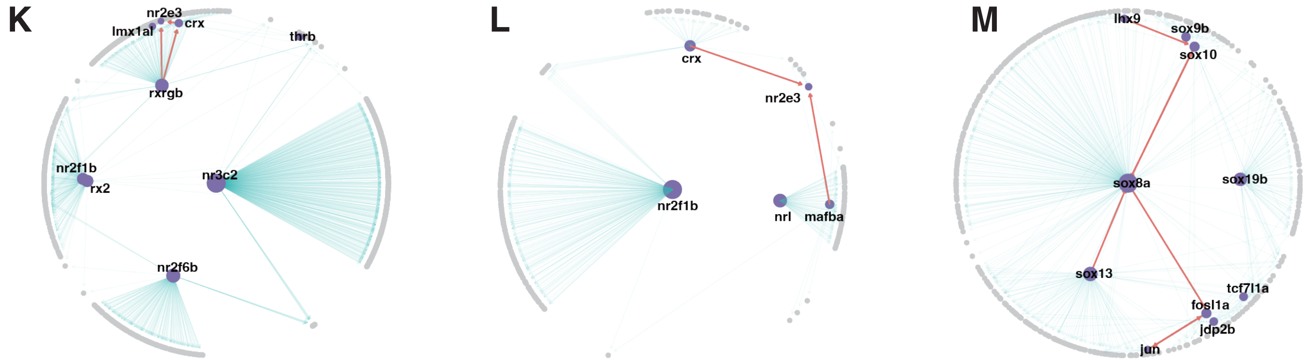


**Figure S5 Reconstruction of GRN in zebrafish retinal cells**

**A–D**: Heatmaps of scaled regulon activity across different stages in zebrafish. (**A**) 24hpf, (**B**) 36hpf, (**C**) 48hpf, (**D**) 14dpf. Each row represents a distinct regulon, while each column denotes a specific cell type. Each regulon is named following the format "TF (n)", where "TF" represents the regulator and "n" denotes the number of target genes in that regulon. Intensity of red signifies high activity, whereas blue indicates low activity. **E–M:** GRN derived from the regulons in **A–D** and Figure **3D**. (**E**) 24hpf module 1. (**F**) 36hpf module 2. (**G**) 36hpf RGC. (**H**) 72hpf AC. (**I**) 72hpf HC, (**J**) 72hpf BC, (**K**) 72hpf Cone, (**L**) 72hpf Rod, (**M**) 72hpf MG. These networks showcase the interplay among genes and regulators within the respective cell types. Purple nodes represent regulators, while gray nodes indicate target genes. Red edges depict regulatory interactions between regulators, and gray edges highlight regulatory relationships of regulators to other genes.

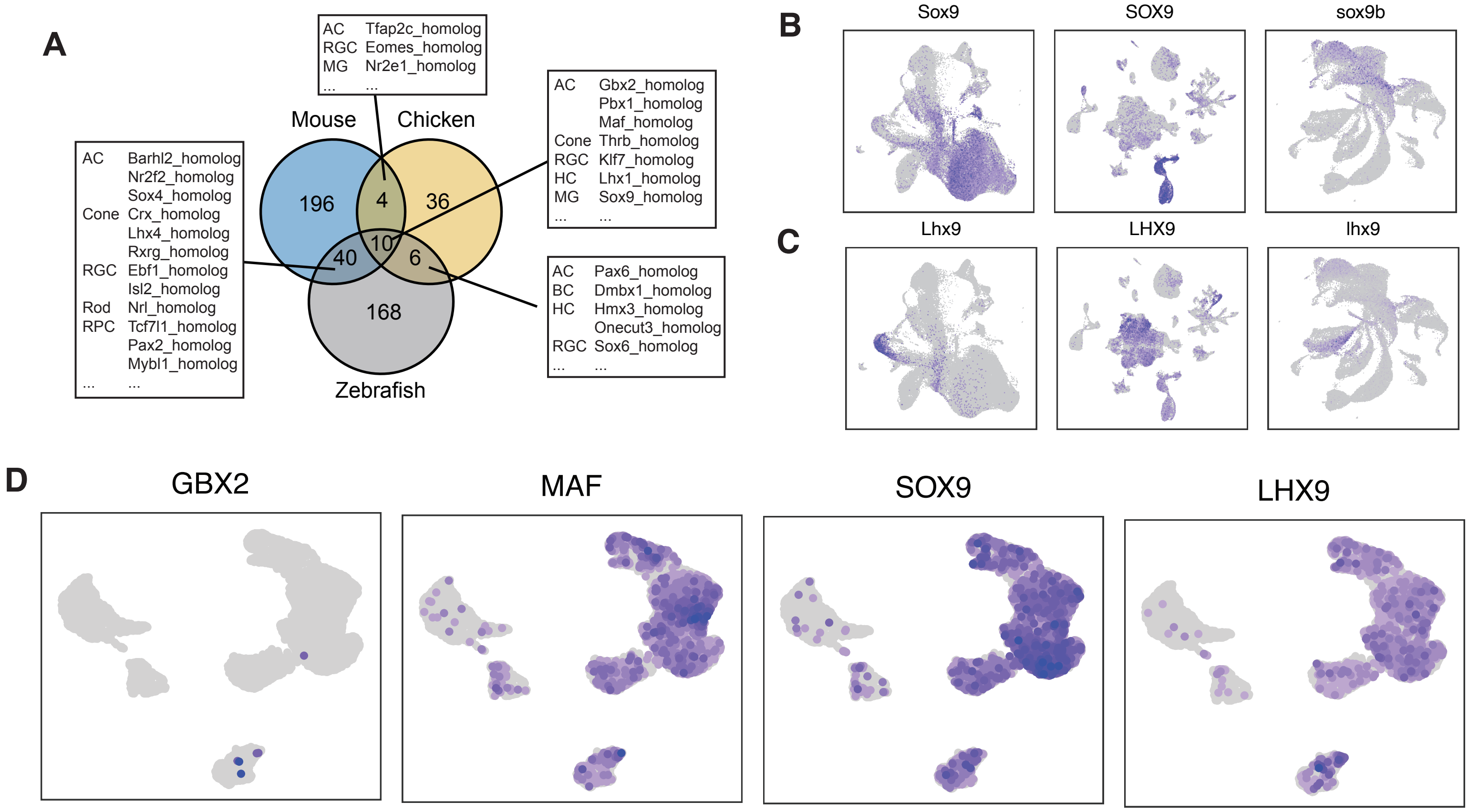


**Figure S6 Expression of cell type specific regulators that are active in all three species**
**A**: Venn diagram illustrating the overlap of regulators in mouse, chicken, and zebrafish. **B-C**: UMAP expression visualizations of (**B**) MG Sox9, (**C**) AC Lhx9 from the shared regulators. Order from left to right: mouse, chicken, zebrafish. Blue indicates high expression; grey indicates low expression. D: UMAP expression visualizations of GBX2, MAF, SOX9 and LHX9 in human retina organoid dataset.

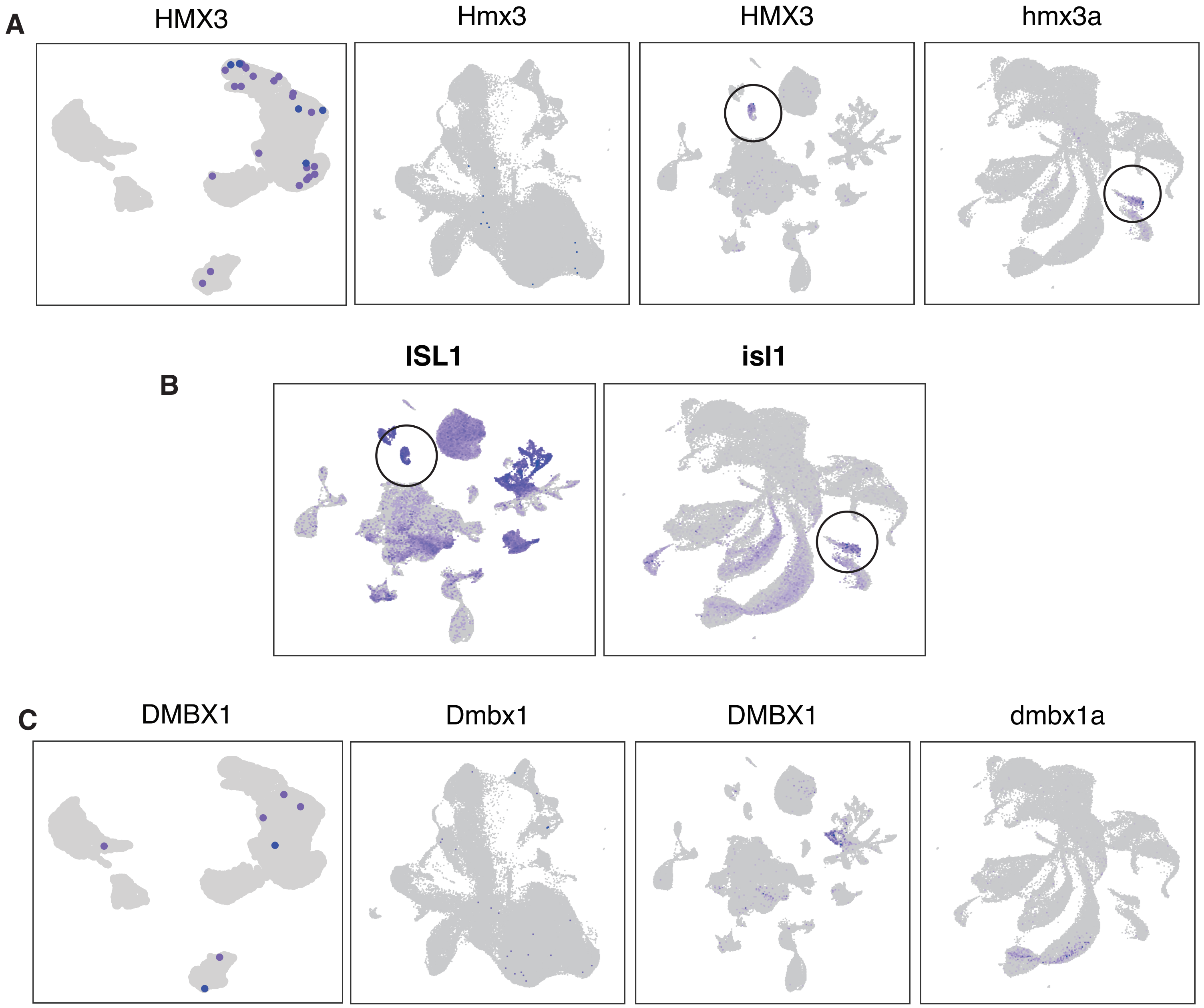


**Figure S7 Expression of Hmx3-homolog and Dmbx1-homolog that are only active in chicken and zebrafish** **A** UMAP expression visualizations of HC-specific regulator Hmx3-homolog. **B** UMAP expression visualizations of Type II HC marker ISL1 (isl1) in chicken and zebrafish. **C** UMAP expression visualizations of HC-specific regulator Hmx3-homolog. Order from left to right: human, mouse, chicken, zebrafish. Blue indicates high expression; grey indicates low expression.

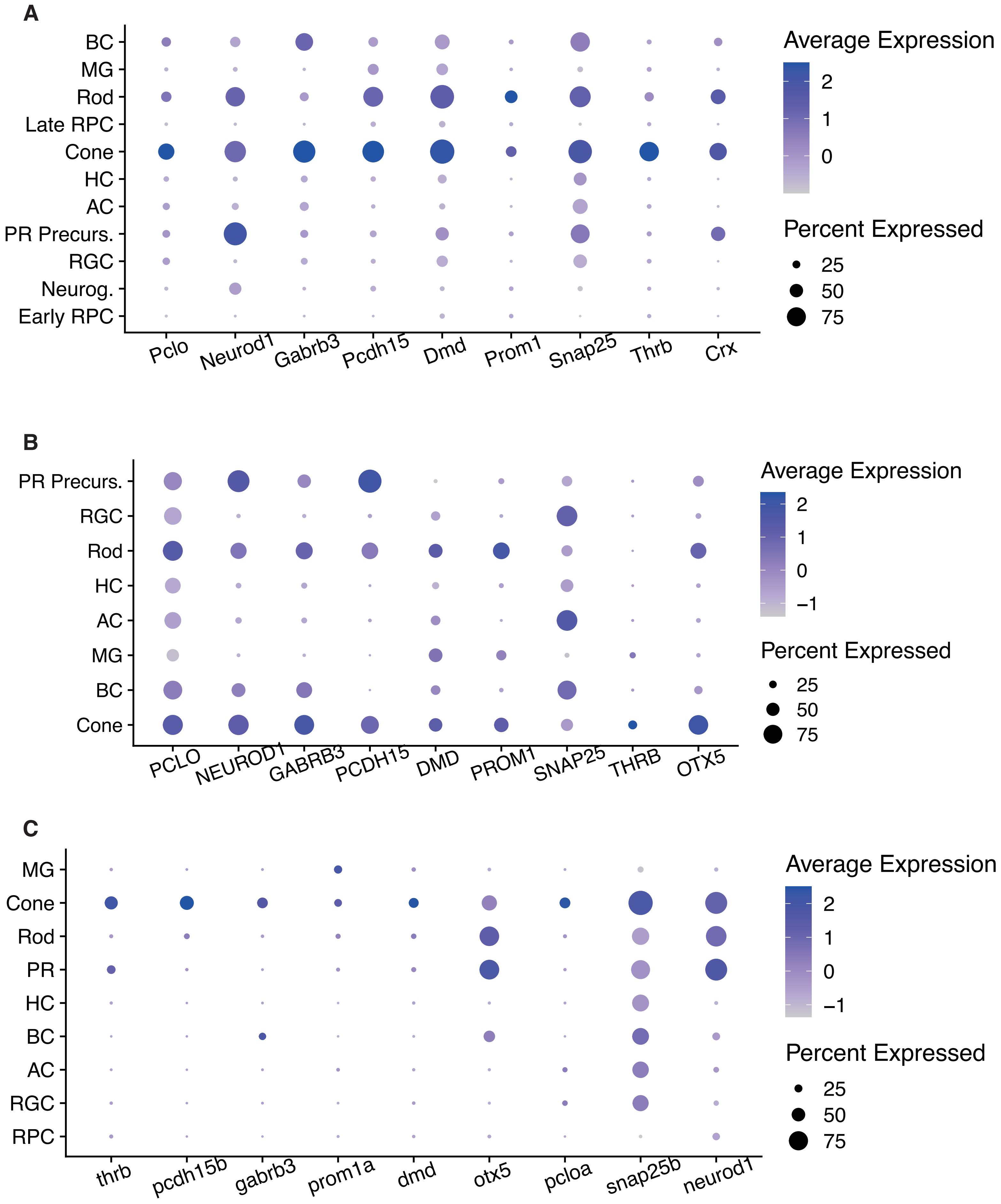


**Figure S8 The expression top genes in E14_Lhx4 regulon that identified as active in mouse cones**
**A–C:** Dotplots showcasing high-expression genes from the E14_Lhx4 regulon across various species. (**A**) mouse, (**B**) chicken, and (**C**) zebrafish

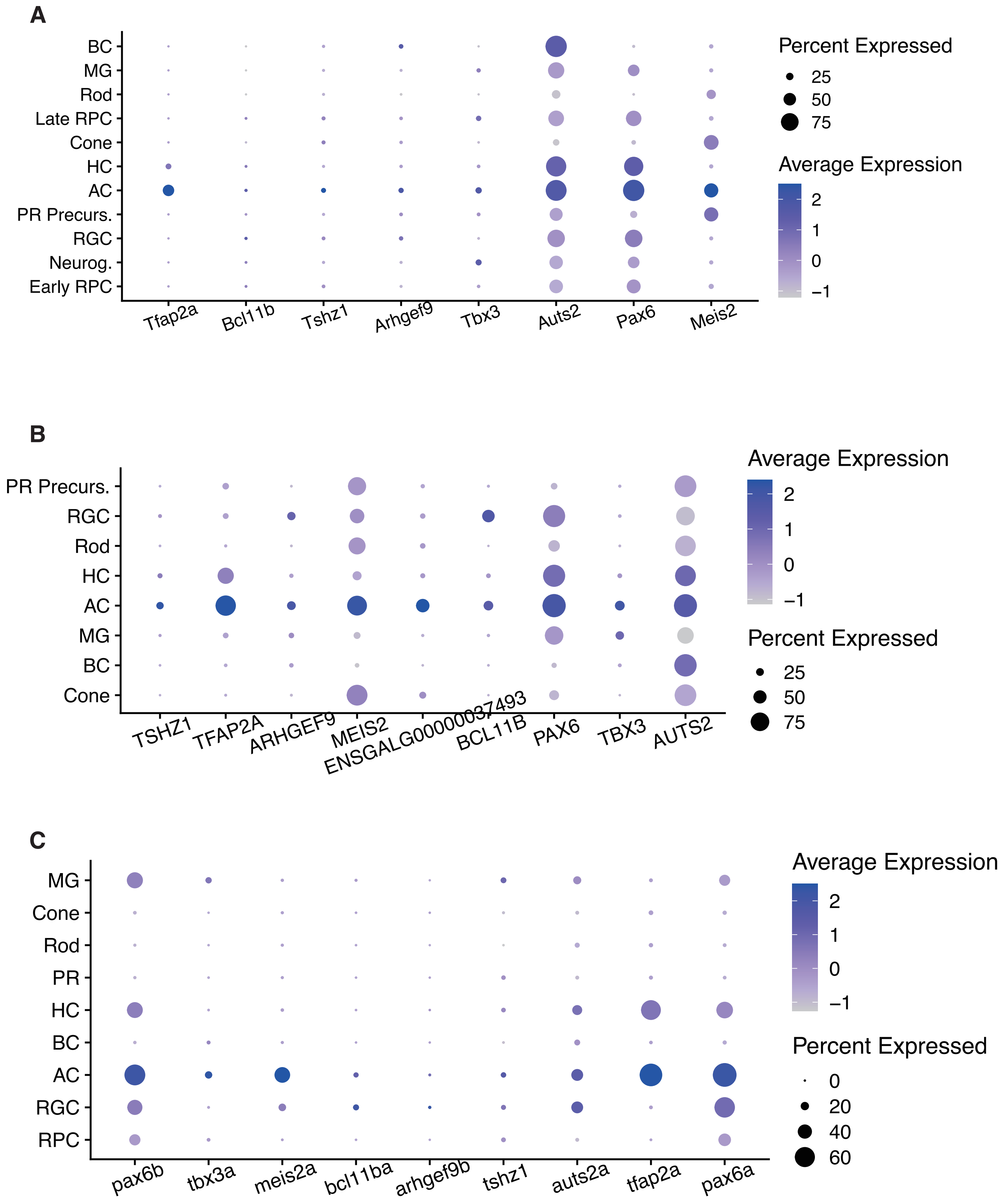


**Figure S9 The expression top genes in E12_PBX1 regulon that identified as active in chicken ACs**
**A–C:** Dotplots showcasing high-expression genes from the E12_PBX1 regulon across various species. (**A**) mouse, (**B**) chicken, and (**C**) zebrafish

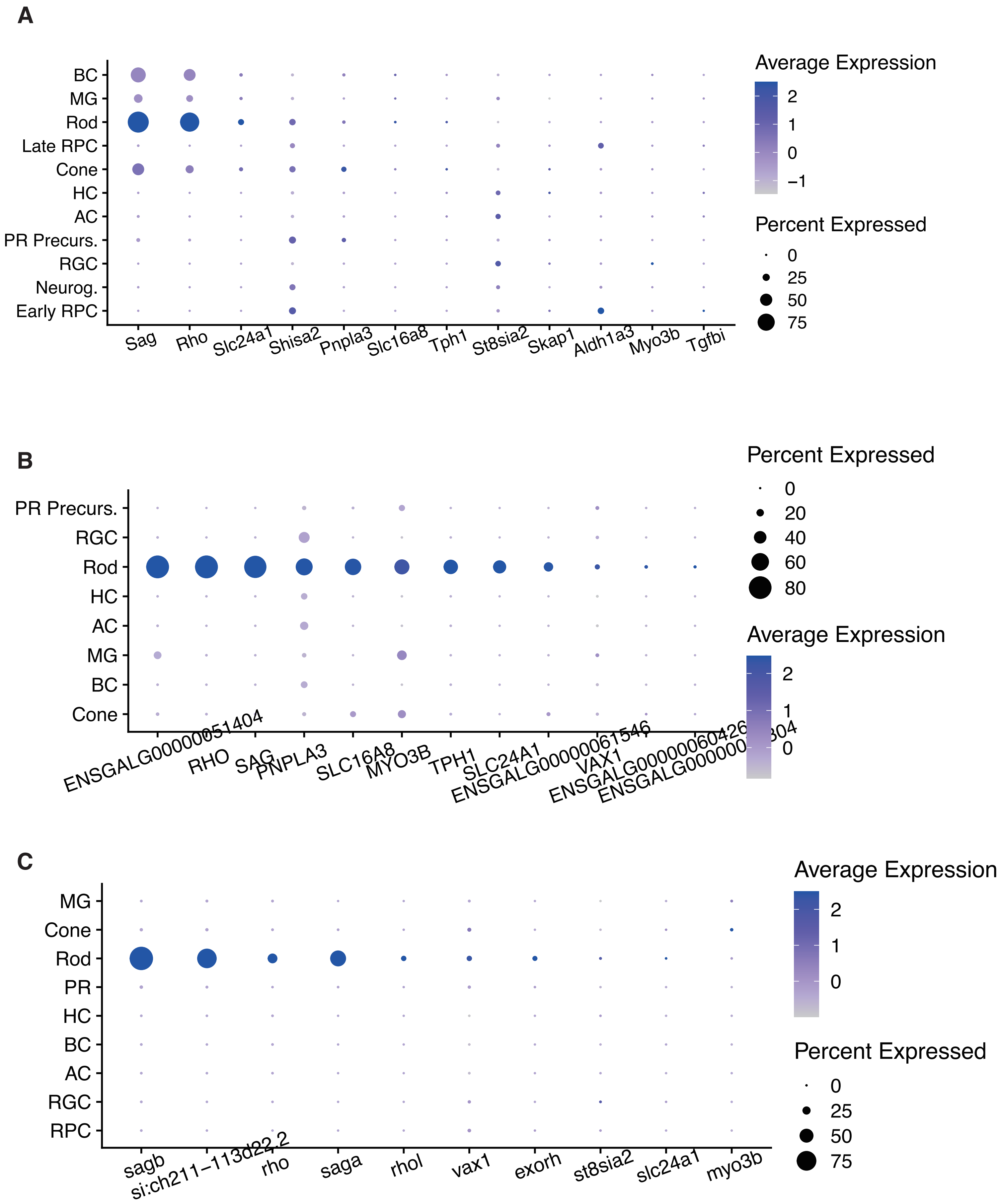


**Figure S10 The expression top genes in E18_VAX1 regulon that identified as active in chicken rods**
**A–C:** Dotplots showcasing high-expression genes from the E18_VAX1 regulon across various species. (**A**) mouse, (**B**) chicken, and (**C**) zebrafish

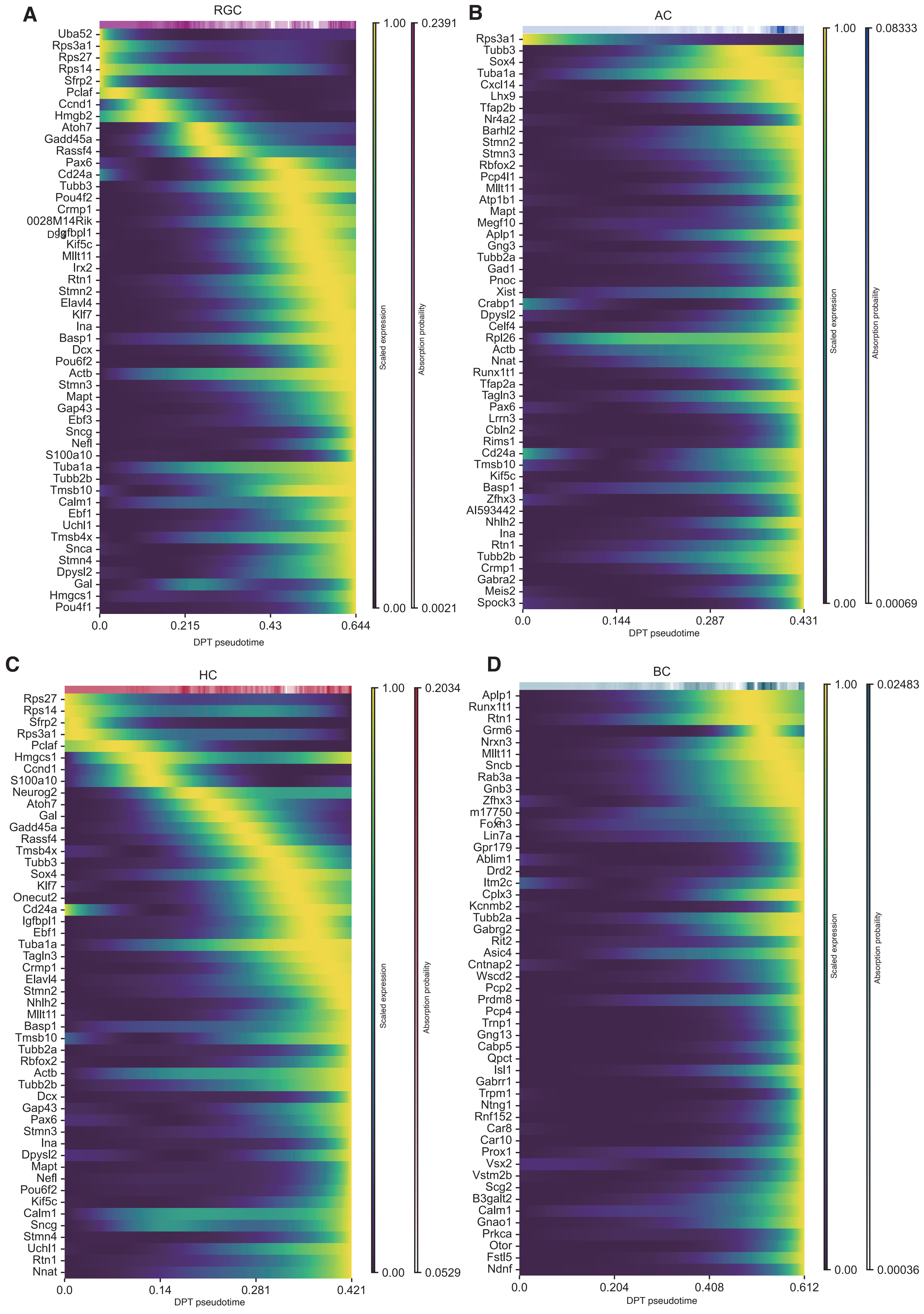

A
RGC
Uba52
Rps3a1
Rps27
Rps14
Sfrp2
Pclaf
Ccnd1
Hmgb2
Atoh7
Gadd45a
Rassf4
Pax6
Cd24a
Tubb3
Pou4f2
Crmp1
0028M14Rik
Igfbpl1
Kif5c
Mllt11
Irx2
Rtn1
Stmn2
Elavl4
Klf7
Ina
Basp1
Dcx
Pou6f2
Actb
Stmn3
Mapt
Gap43
Ebf3
Sncg
Nefl
S100a10
Tuba1a
Tubb2b
Tmsb10
Calm1
Ebf1
Uchl1
Tmsb4x
Snca
Stmn4
Dpysl2
Gal
Hmgcs1
Pou4f1
0.0
0.215
0.43
0.644
DPT pseudotime
1.00
0.00
Scaled expression
0.2391
0.0021
Absorption probability
B
AC
Rps3a1
Tubb3
Sox4
Tuba1a
Cxcl14
Lhx9
Tfap2b
Nr4a2
Barhl2
Stmn2
Stmn3
Rbfox2
Pcp4l1
Mllt11
Atp1b1
Mapt
Megf10
Aplp1
Gng3
Tubb2a
Gad1
Pnoc
Xist
Crabp1
Dpysl2
Celf4
Rpl26
Actb
Nnat
Runx1t1
Tfap2a
Tagln3
Pax6
Lrrn3
Cbln2
Rims1
Cd24a
Tmsb10
Kif5c
Basp1
Zfhx3
AI593442
Nhlh2
Ina
Rtn1
Tubb2b
Crmp1
Gabra2
Meis2
Spock3
0.0
0.144
0.287
0.431
DPT pseudotime
1.00
0.00
Scaled expression
0.08333
0.00069
Absorption probability
C
HC
Rps27
Rps14
Sfrp2
Rps3a1
Pclaf
Hmgcs1
Ccnd1
S100a10
Neurog2
Atoh7
Gal
Gadd45a
Rassf4
Tmsb4x
Tubb3
Sox4
Klf7
Onecut2
Cd24a
Igfbpl1
Ebf1
Tuba1a
Tagln3
Crmp1
Elavl4
Stmn2
Nhlh2
Mllt11
Basp1
Tmsb10
Tubb2a
Rbfox2
Actb
Tubb2b
Dcx
Gap43
Pax6
Stmn3
Ina
Dpysl2
Mapt
Nefl
Pou6f2
Kif5c
Calm1
Sncg
Stmn4
Uchl1
Rtn1
Nnat
0.0
0.14
0.281
0.421
DPT pseudotime
1.00
0.00
Scaled expression
0.2034
0.0529
Absorption probaility
D
BC
Aplp1
Runx1t1
Rtn1
Grm6
Nrxn3
Mllt11
Sncb
Rab3a
Gnb3
Zfhx3
m17750
Foxn3
Lin7a
Gpr179
Ablim1
Drd2
Itm2c
Cplx3
Kcnmb2
Tubb2a
Gabrg2
Rit2
Asic4
Cntnap2
Wscd2
Pcp2
Prdm8
Pcp4
Trnp1
Gng13
Cabp5
Qpct
Isl1
Gabrr1
Trpm1
Ntng1
Rnf152
Car8
Car10
Prox1
Vsx2
Vstm2b
Scg2
B3galt2
Calm1
Gnao1
Prkca
Otor
Fstl5
Ndnf
0.0
0.204
0.408
0.612
DPT pseudotime
1.00
0.00
Scaled expression
0.02483
0.00036
Absorption probaility

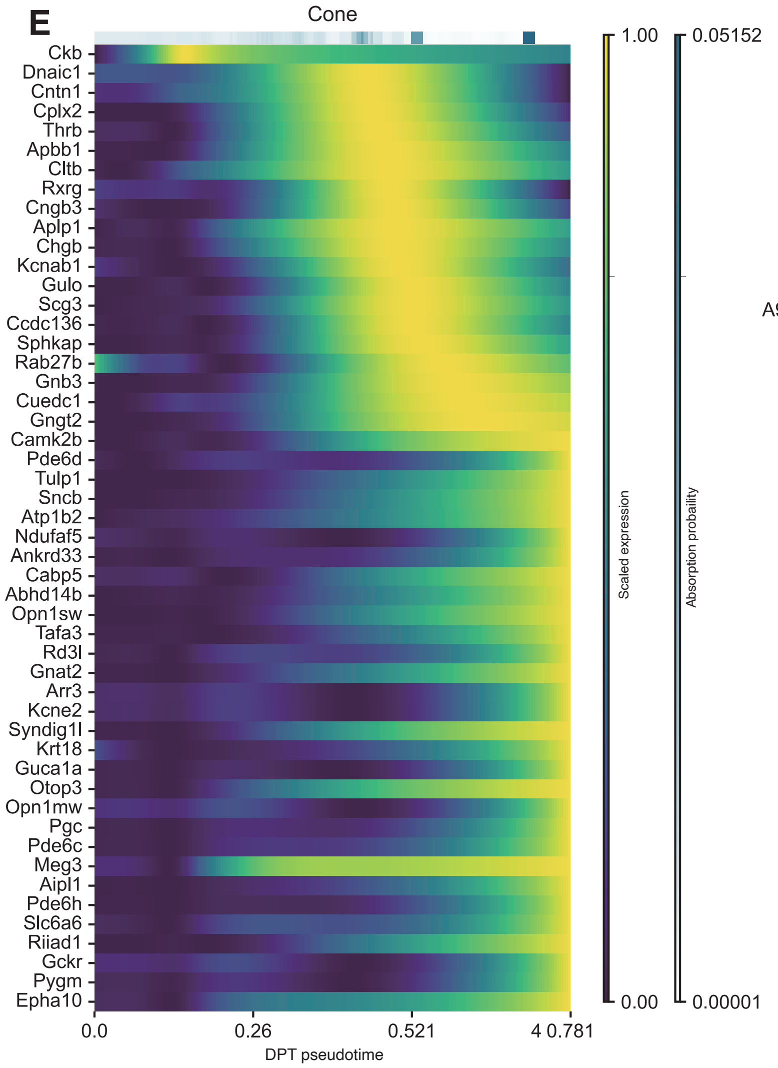


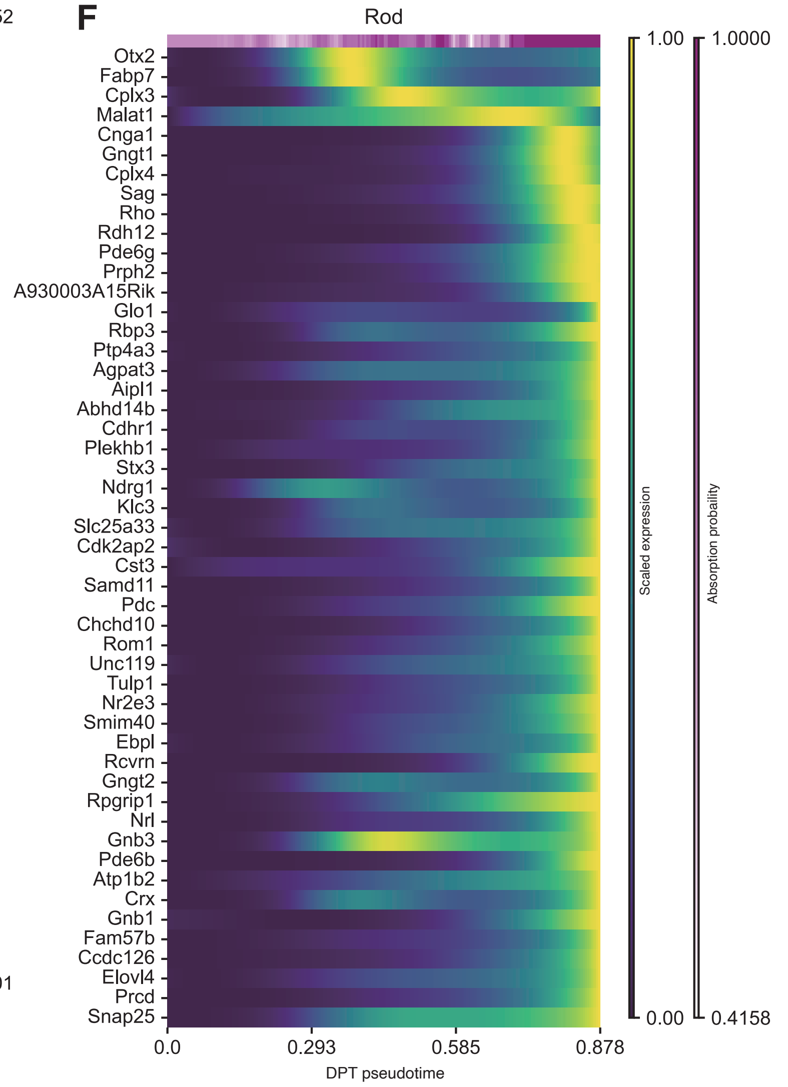


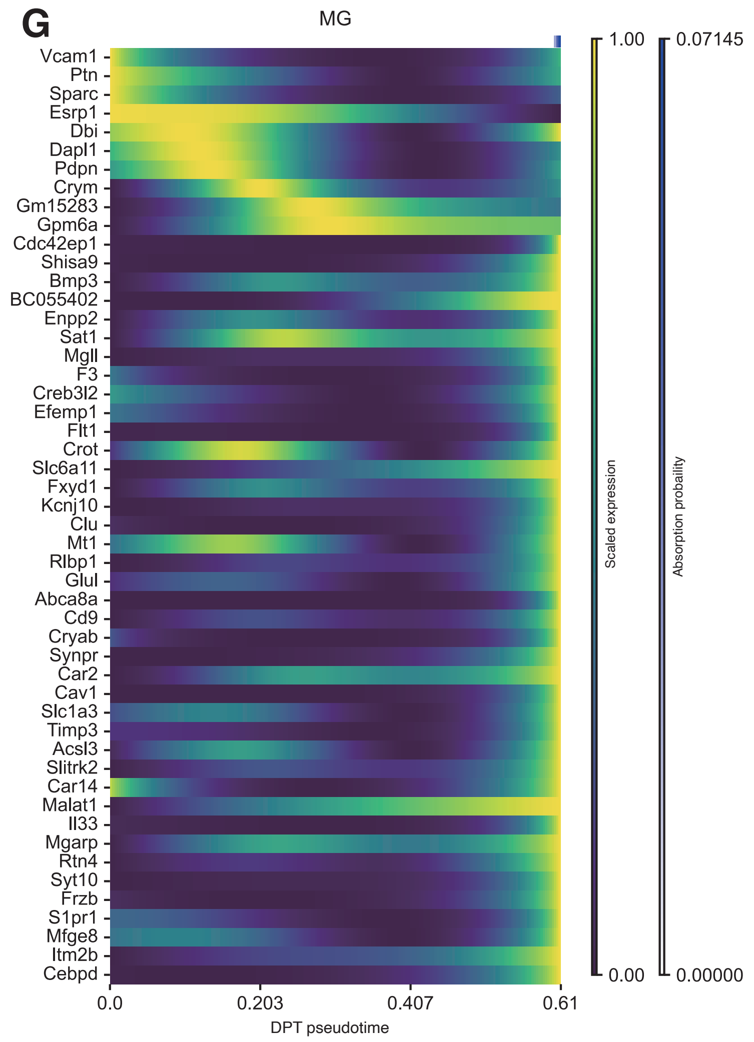


**Figure S11 Gene expression trends of putative driver genes along DPT pesudotime in mouse retina.**
Heatmaps illustrating the expression levels of the top 50 cell class-specific putative driver genes across DPT-pseudotime for (**A**) RGCs, (**B**) ACs, (**C**) HCs, (**D**) BCs, (**E**) Cones, (**F**) Rods, (**G**) MG

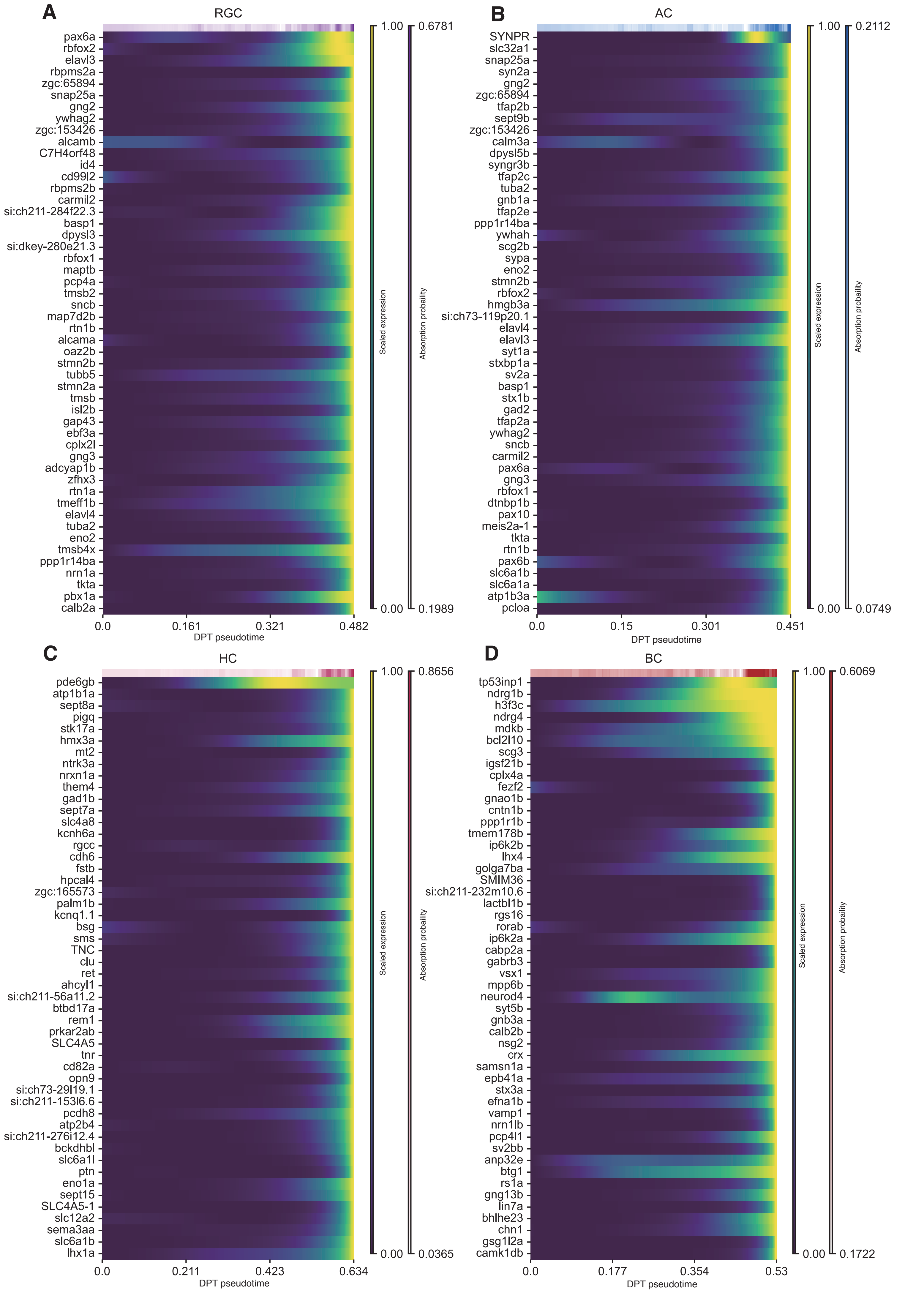
A
RGC
pax6a
rbfox2
elavl3
rbpms2a
zgc:65894
snap25a
gng2
ywhag2
zgc:153426
alcamb
C7H4orf48
id4
cd99l2
rbpms2b
carmil2
si:ch211-284f22.3
basp1
dpysl3
si:dkey-280e21.3
rbfox1
maptb
pcp4a
tmsb2
sncb
map7d2b
rtn1b
alcama
oaz2b
stmn2b
tubb5
stmn2a
tmsb
isl2b
gap43
ebf3a
cplx2l
gng3
adcyap1b
zfhx3
rtn1a
tmeff1b
elavl4
tuba2
eno2
tmsb4x
ppp1r14ba
nrn1a
tkta
pbx1a
calb2a
0.0
0.161
0.321
0.482
DPT pseudotime
1.00
0.00
Scaled expression
0.6781
0.1989
Absorption probability
B
AC
SYNPR
slc32a1
snap25a
syn2a
gng2
zgc:65894
tfap2b
sept9b
zgc:153426
calm3a
dpysl5b
syngr3b
tfap2c
tuba2
gnb1a
tfap2e
ppp1r14ba
ywhah
scg2b
sypa
eno2
stmn2b
rbfox2
hmgb3a
si:ch73-119p20.1
elavl4
elavl3
syt1a
stxbp1a
sv2a
basp1
stx1b
gad2
tfap2a
ywhag2
sncb
carmil2
pax6a
gng3
rbfox1
dtnbp1b
pax10
meis2a-1
tkta
rtn1b
pax6b
slc6a1b
slc6a1a
atp1b3a
pcloa
0.0
0.15
0.301
0.451
DPT pseudotime
1.00
0.00
Scaled expression
0.2112
0.0749
Absorption probability
C
HC
pde6gb
atp1b1a
sept8a
pigq
stk17a
hmx3a
mt2
ntrk3a
nrxn1a
them4
gad1b
sept7a
slc4a8
kcnh6a
rgcc
cdh6
fstb
hpcal4
zgc:165573
palm1b
kcnq1.1
bsg
sms
TNC
clu
ret
ahcyl1
si:ch211-56a11.2
btbd17a
rem1
prkar2ab
SLC4A5
tnr
cd82a
opn9
si:ch73-29l19.1
si:ch211-153l6.6
pcdh8
atp2b4
si:ch211-276i12.4
bckdhbl
slc6a1l
ptn
eno1a
sept15
SLC4A5-1
slc12a2
sema3aa
slc6a1b
lhx1a
0.0
0.211
0.423
0.634
DPT pseudotime
1.00
0.00
Scaled expression
0.8656
0.0365
Absorption probability
D
BC
tp53inp1
ndrg1b
h3f3c
ndrg4
mdkb
bcl2l10
scg3
igsf21b
cplx4a
fezf2
gnao1b
cntn1b
ppp1r1b
tmem178b
ip6k2b
lhx4
golga7ba
SMIM36
si:ch211-232m10.6
lactbl1b
rgs16
rorab
ip6k2a
cabp2a
gabrb3
vsx1
mpp6b
neurod4
syt5b
gnb3a
calb2b
nsg2
crx
samsn1a
epb41a
stx3a
efna1b
vamp1
nrn1lb
pcp4l1
sv2bb
anp32e
btg1
rs1a
gng13b
lin7a
bhlhe23
chn1
gsg1l2a
camk1db
0.0
0.177
0.354
0.53
DPT pseudotime
1.00
0.00
Scaled expression
0.6069
0.1722
Absorption probaility

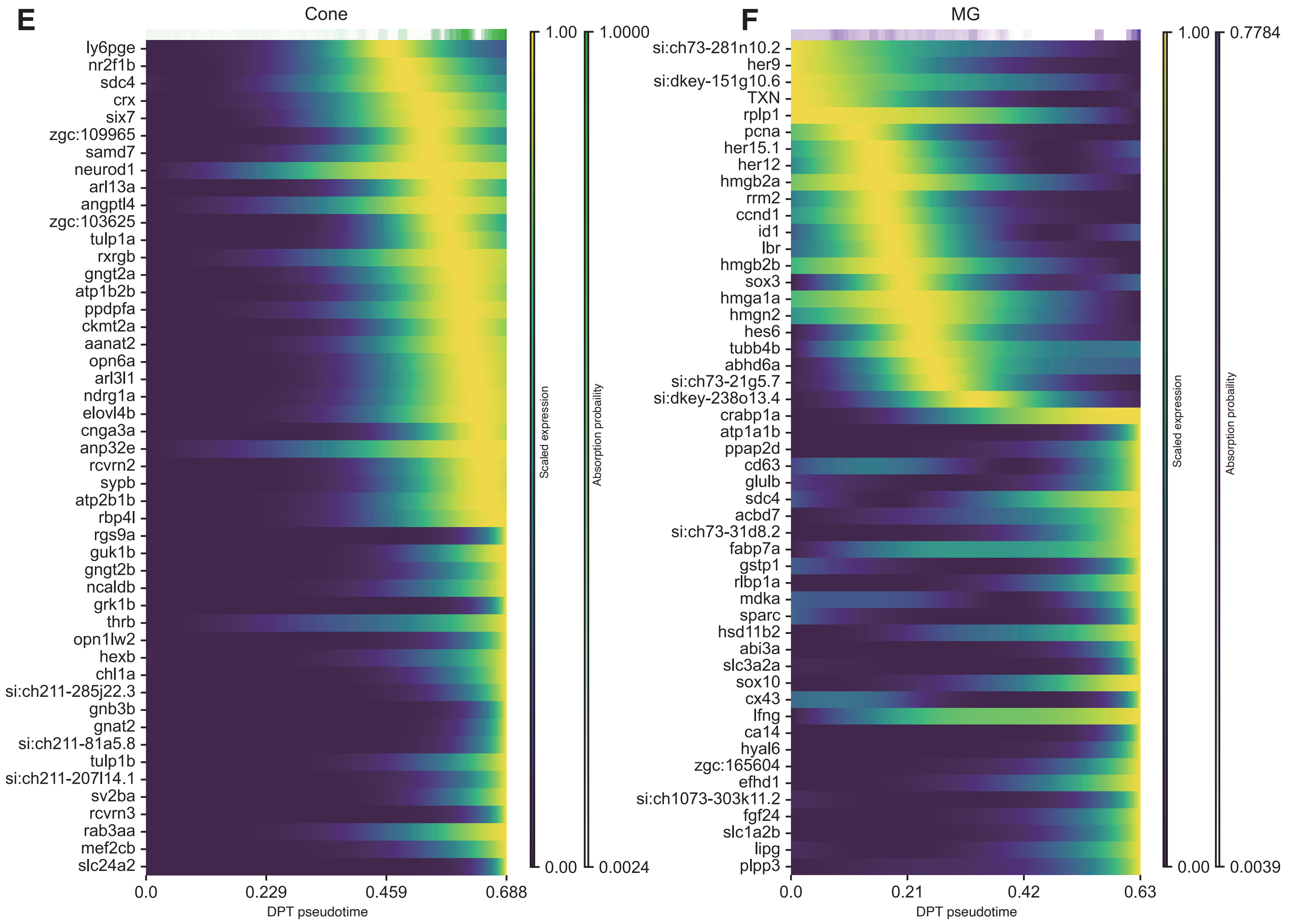


**Figure S12 Gene expression trends of putative driver genes along DPT pesudotime in zebrtafish retina.** Heatmaps illustrating the expression levels of the top 50 cell class-specific putative driver genes across DPT-pseudotime for (**A**) RGCs, (**B**) ACs, (**C**) HCs, (**D**) BCs, (**E**) Cones, (**F**) MG.

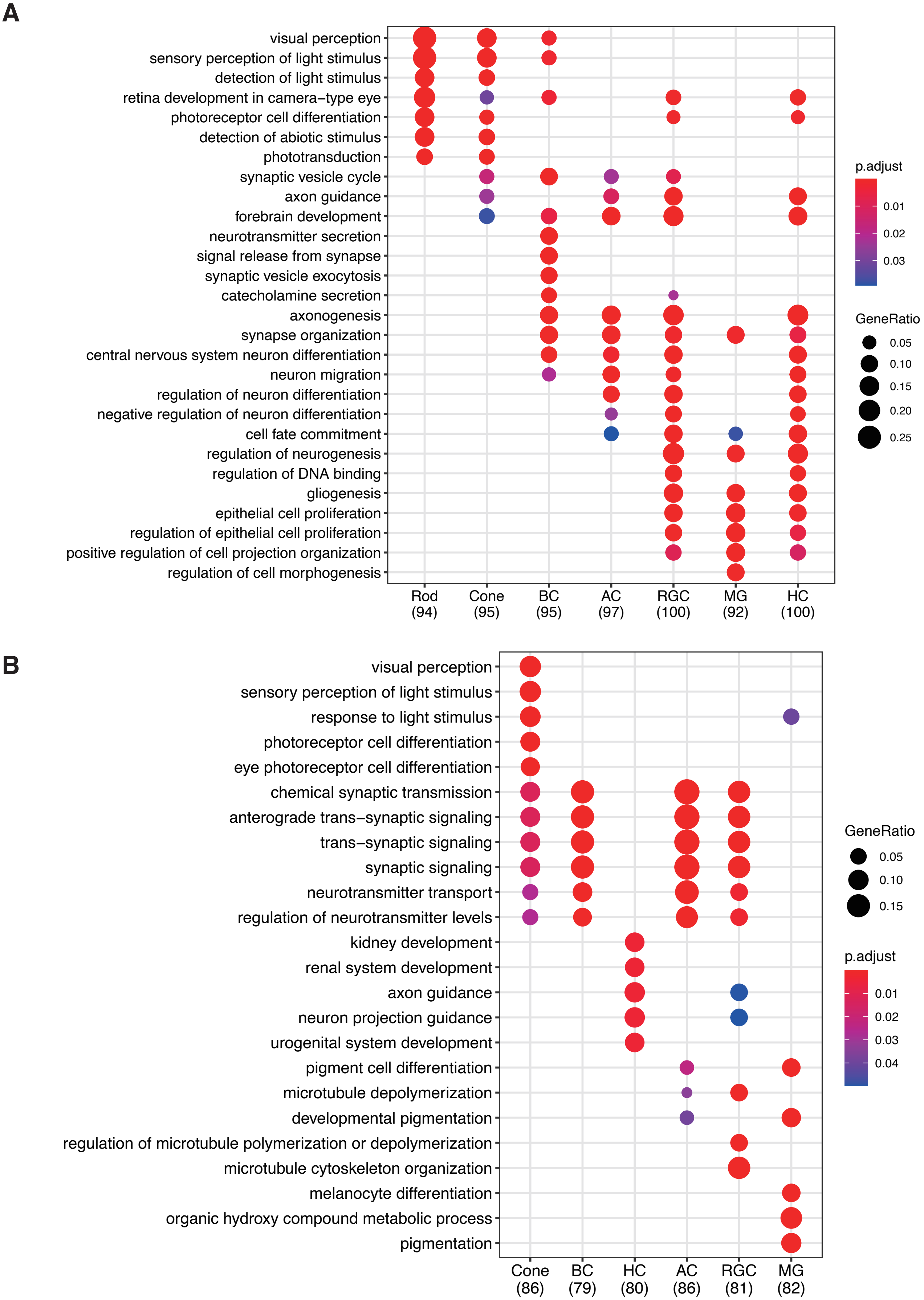


**Figure S13 Gene Ontology (GO) analysis of cell class-specific putative driver genes in mouse and zebrafish**

Dotplots of GO analysis in biological process for the top 100 cell class-specific putative driver genes. (**A**) mouse. (**B**) zebrafish

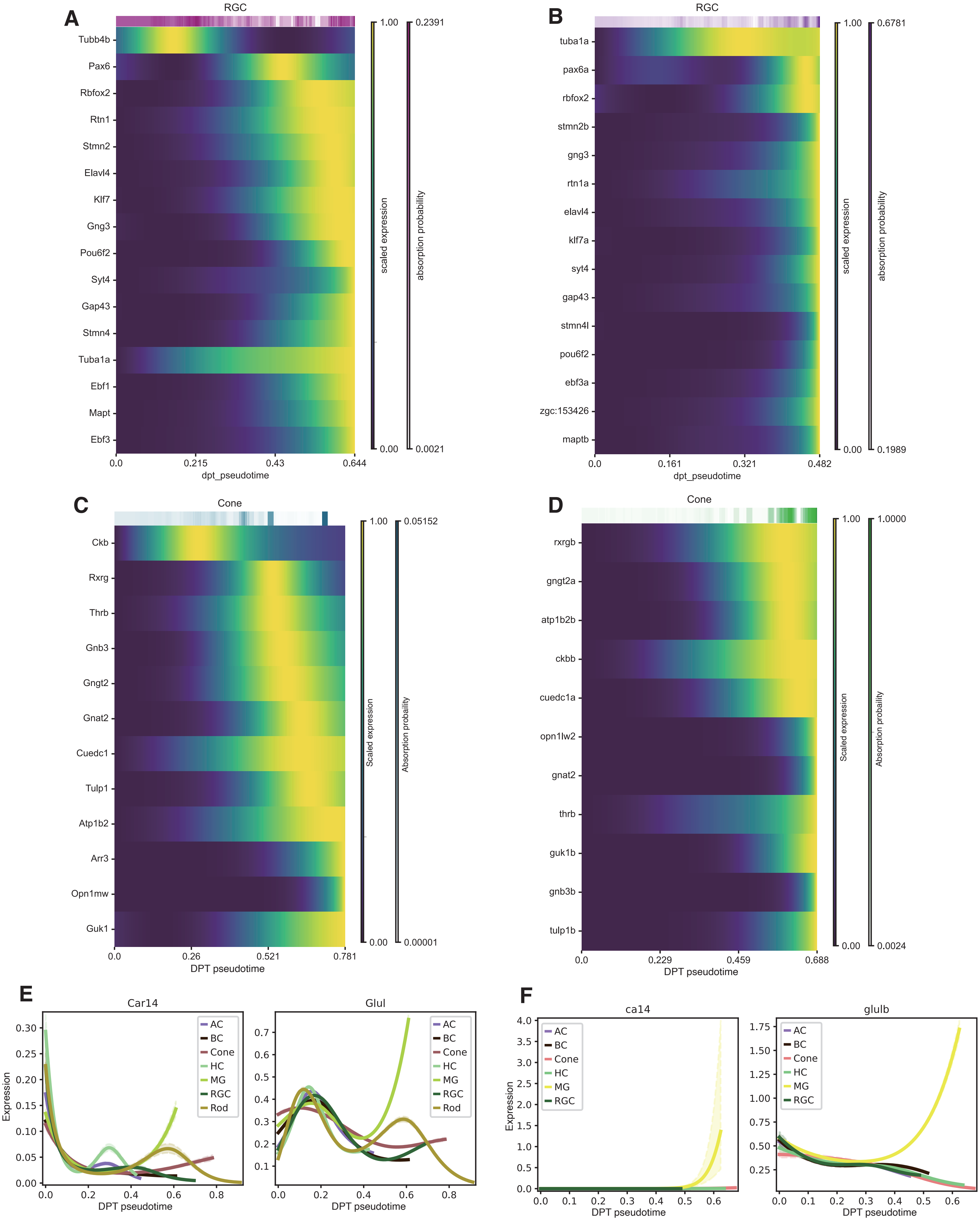


**Figure S14 Gene expression trends of shared driver genes along DPT pesudotime between mouse and zebrafish.**

Heatmaps illustrating the expression levels of shared driver genes in mouse ACs (**A**), zebrafish ACs (**B**), mouse Cones (**C**), zebrafish Cones (**D**). mouse MG (**E**), and zebrafish MG (**F**) across DPT-pseudotime.

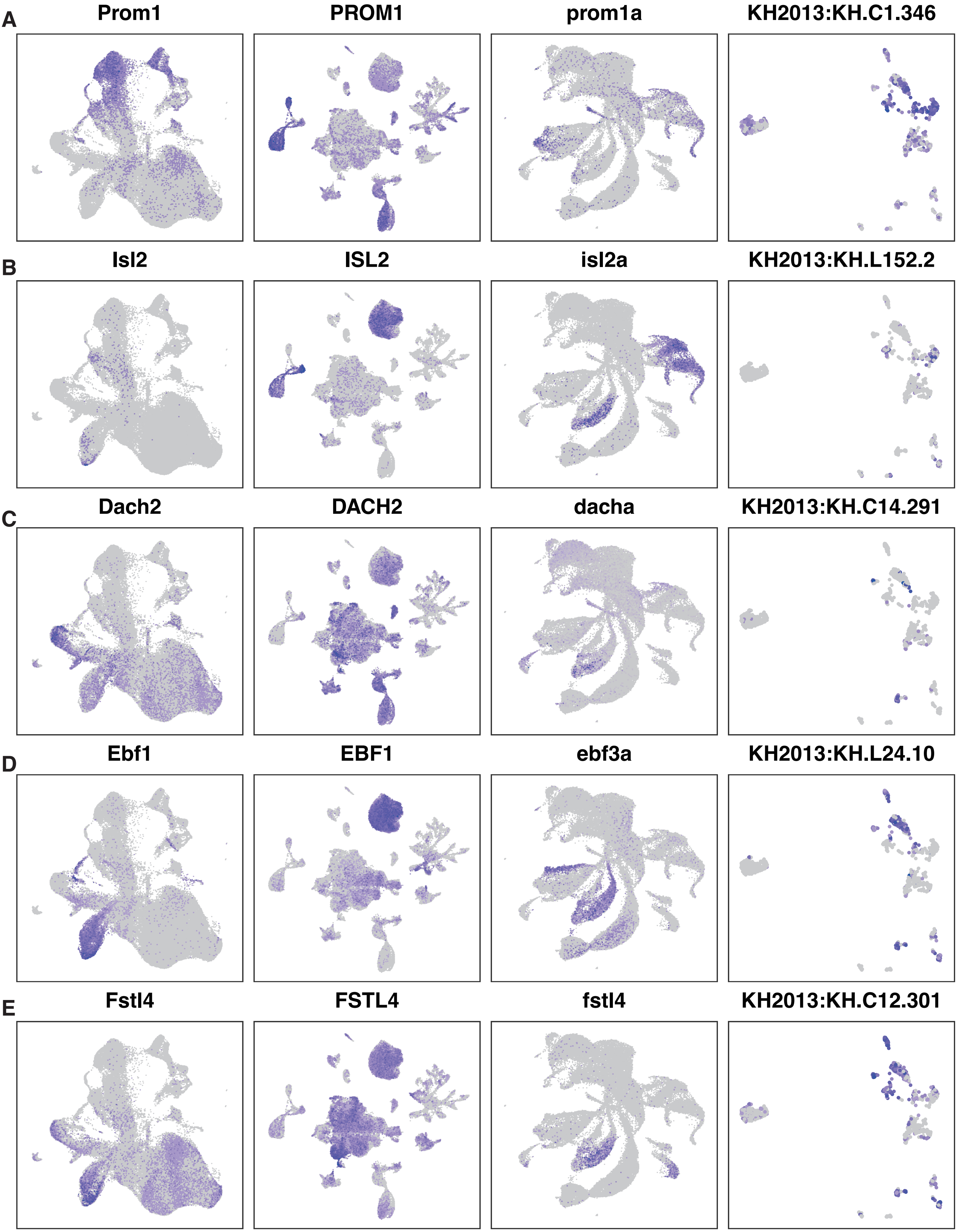
A
Prom1
PROM1
prom1a
KH2013:KH.C1.346
B
Isl2
ISL2
isl2a
KH2013:KH.L152.2
C
Dach2
DACH2
dacha
KH2013:KH.C14.291
D
Ebf1
EBF1
ebf3a
KH2013:KH.L24.10
E
Fstl4
FSTL4
fstl4
KH2013:KH.C12.301

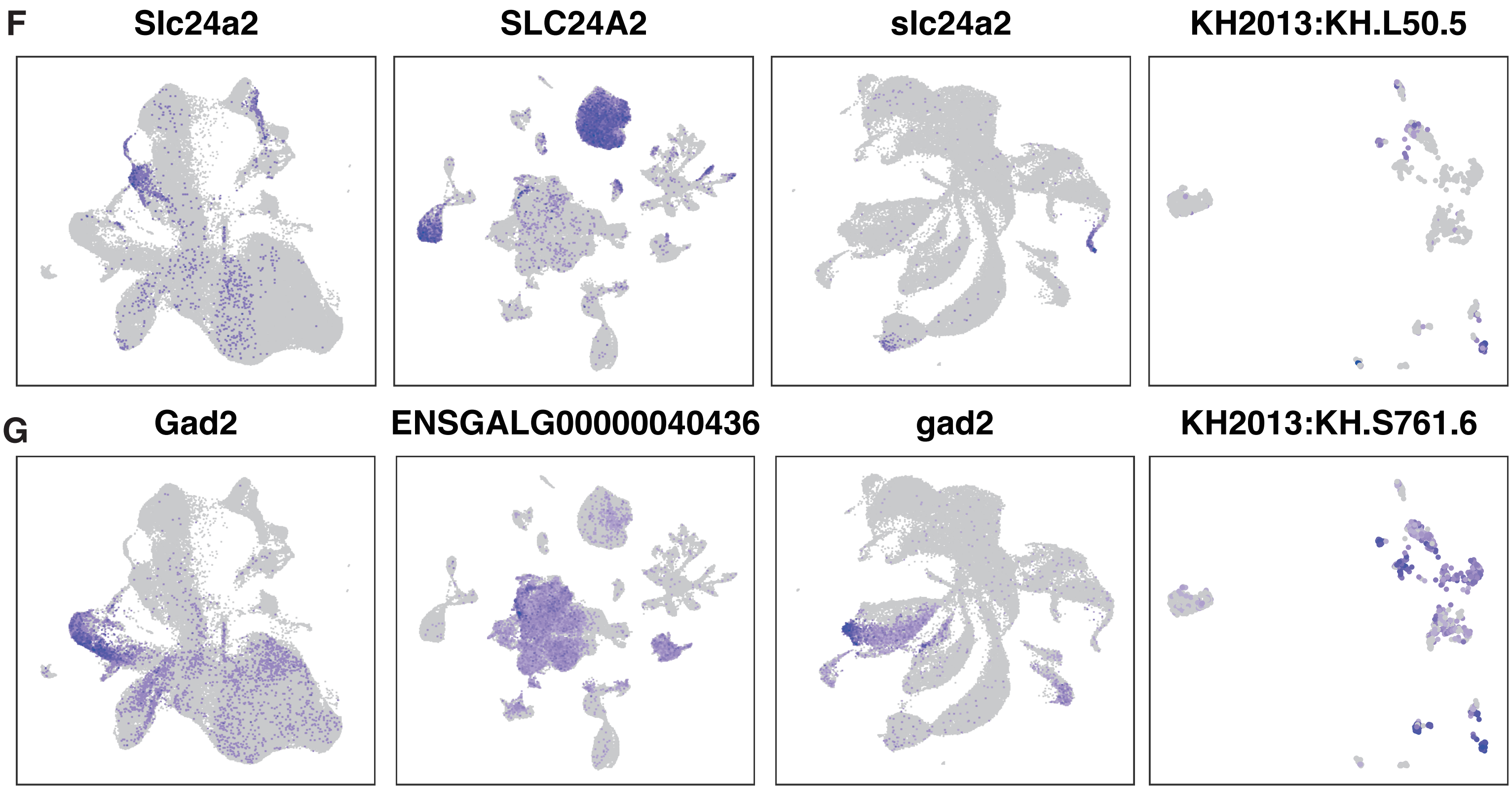


**Figure S15 Shared DEGs between vertebrate retinal cells and *Ciona* PR-related cells.**
UMAP visualizations of gene expression levels across four species. Each figure represents the expression of a specific gene: (**A**) Prom1, (**B**) Isl2, (**C**) Dach2, (**D**) Ebf1,(**E**) Fstl4,(**F**) Slc24a, (**G**) Gad2. From left to right, species are mouse, chicken, zebrafish, and *Ciona*. Blue indicates high expression, gray indicates low expression.

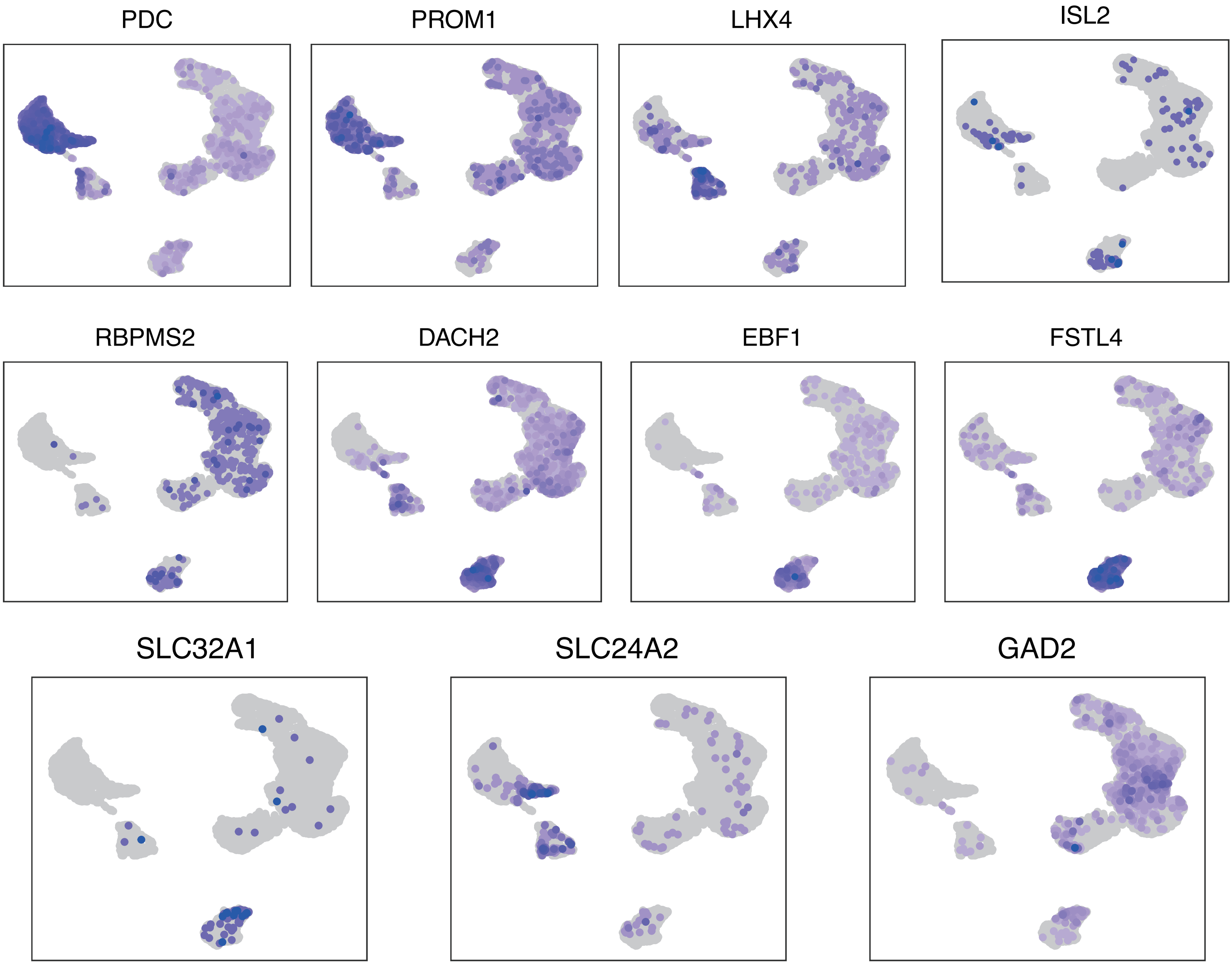


**Figure S16 Expression of genes in human retina organoid development dataset.**
**A–E**: UMAP visualizations of (**A**) AC Tfap2c-homolog, (**B**) RGC Eomes-homolog (**C**) AC Pax6-homolog, (**D**) RGC Sox6-homolog, (**E**) HC Onecut3-homolog from the shared regulators in Figure . Order from left to right: mouse, chicken, zebrafish. Blue indicates high expression; grey indicates low expression.
